# scifAI: Explainable machine learning for profiling the immunological synapse and functional characterization of therapeutic antibodies

**DOI:** 10.1101/2022.10.24.513494

**Authors:** Sayedali Shetab Boushehri, Katharina Essig, Nikolaos-Kosmas Chlis, Sylvia Herter, Marina Bacac, Fabian J Theis, Elke Glasmacher, Carsten Marr, Fabian Schmich

**Affiliations:** Institute of AI for Health, Helmholtz Zentrum München, German Research Center for Environmental Health, Neuherberg, Germany; Institute of Computational Health, Helmholtz Zentrum München, German Research Center for Environmental Health, Neuherberg, Germany; Technical University of Munich, Department of Mathematics, Munich, Germany; Data Science, Pharmaceutical Research and Early Development Informatics (pREDi), Roche Innovation Center Munich, Germany; Large Molecule Research (LMR), Roche Pharmaceutical Research and Early Development (pRED), Roche Innovation Center Munich, Germany; Discovery Oncology, Roche Pharmaceutical Research and Early Development (pRED), Roche Innovation Zurich, Switzerland; RED (Research and Early Development), Roche Diagnostics Solutions, Roche Innovation Center Munich, Germany

**Keywords:** immunological synapse, mechanistic insight, therapeutic antibodies, imaging flow cytometry, IFC, functional characterization, explainable, machine learning, AI, feature extraction, prediction, classification, computer vision, deep learning, python, framework

## Abstract

Therapeutic antibodies are widely used to treat severe diseases. Most of them alter immune cells and act within the immunological synapse; an essential cell-to-cell interaction to direct the humoral immune response. Although many antibody designs are generated and evaluated, a high-throughput tool for systematic antibody characterization and prediction of function is lacking. Here, we introduce the first comprehensive open-source framework, scifAI (single-cell imaging flow cytometry AI), for preprocessing, feature engineering and explainable, predictive machine learning on imaging flow cytometry (IFC) data. Additionally, we generate the largest publicly available IFC data set of the human immunological synapse containing over 2.8 million images. Using scifAI, we analyze class frequency- and morphological changes under different immune stimulation. T cell cytokine production across multiple donors and therapeutic antibodies is quantitatively predicted *in vitro,* linking morphological features with function and demonstrating the potential to significantly impact antibody design. scifAI is universally applicable to IFC data. Given its modular architecture it is straightforward to incorporate into existing workflows and analysis pipelines, e.g. for rapid antibody screening and functional characterization.

## Introduction

The formation of an immunological synapse is the first event of the adaptive immune reaction induced by the interaction of a T cell with its corresponding antigen-presenting cell (APC). This rapidly formed cell-cell interface is initiated by the recognition of peptide-loaded major histocompatibility complexes (MHC) by the T cell receptor (TCR). It involves the rearrangement of actin filaments of the cytoskeleton and the recruitment of signaling, co-stimulatory, co-inhibitory, and adhesion molecules to the nascent synapse ^1, 2^. This process is crucial to trigger and fine-tune T cell responses and ensure intact immune reactions. Dysfunctional immunological synapse formation has been observed in several immune-related disorders ^3–8^ and has thus been considered a potential target to stimulate or inhibit immune responses by modulating its assembly or function ^9–11^. For instance, various therapeutic antibodies were developed that alter immunological synapse formation to treat cancer and autoimmune diseases ^12–15^. Although significant progress in developing immunological synapse targeting agents has been achieved in the last years ^9^, there is still need to refine the compounds further, especially to improve their efficacy. It has been identified that antibody size and format ^16, 17^, the dose, as well as target expression ^18^ can be critical parameters for immunological synapse formation and its effect on T cell function.

However, so far no study has provided a tool to systematically quantify and characterize the morphology of the immunological synapse, investigate its correlation to T cell response, or identify properties predictive for the efficacy of antibodies *in vitro*. As a consequence, only a literature-guided set of fluorescent stainings relevant for investigating the immunological synapse is set in an otherwise untargeted approach, allowing the exploration of a broad range of possible characteristics. The key technology for high-throughput data acquisition for this purpose is imaging flow cytometry (IFC), combining the benefits of traditional flow cytometry with deep, multi-channel imaging on the single-cell level. IFC has recently been successfully applied to visualize and quantify the immunological synapse of primary human T:APC cell conjugates ^19–21^, however, none of these studies investigated the formation of the immunological synapse in the context of T cell effector function (cytokine production).

Recent studies have demonstrated the potential of machine learning algorithms for a more robust and accurate analysis of high-throughput imaging data, an approach that has been demonstrated to overcome limitations of conventional gating strategies ^22–24^. Leveraging machine learning for IFC data analysis has also enabled the identification of morphological patterns in the cell, a combined analysis of RNA and protein data, and the implementation of predictive models ^22–26^. While limited open-source software implementations designed for IFC data analysis are available ^26, 27^, they either rely on additional third-party software adding complexity in the analysis pipeline, or they focus on prediction performance only and lack explainability.

Here, we present scifAI, a machine learning framework for the efficient and explainable analysis of high-throughput imaging data based on a modular open-source implementation. We also publish the largest publicly available multi-channel IFC data set with over 2.8 million images of primary human T-B cell conjugates from multiple donors, and demonstrate how scifAI can be used to detect patterns and build predictive models. We showcase the potential of our framework for (i) the prediction of immunologically relevant cell class frequencies, (ii) the systematic morphological profiling of the immunological synapse, (iii) the investigation of inter donor and inter and intra-experiment variability, as well as (iv) the characterization of the mode of action of therapeutic antibodies and (v) the prediction of their functionality *in vitro.* Combining high-throughput imaging of the immunological synapse using IFC with rigorous data preprocessing and machine learning enables researchers in pharma to screen for novel antibody candidates and improved evaluation of lead molecules in terms of functionality, mode-of-action insights and antibody characteristics such as affinity, avidity and format.

## Results

### Comprehensive multi-channel imaging flow cytometry data set of the immunological synapse

Formation of T cell immune synapses occurs at variable and relatively low frequencies depending on the donor, the APC, and pharmacological perturbations (^9, 12, 19–21)^. Therefore, high-throughput IFC was selected as the method of choice to capture a large number of samples, enabling the detection of subtle changes in cell morphology. Using IFC we generated a comprehensive data set for the systematic analysis of the immunological synapse of T-B conjugates (Fig. 1a and Supplementary Fig. 1a). Human memory CD4^+^ T cells, isolated from peripheral blood of different donors, were co-cultured with superantigen (*Staphylococcus aureus* enterotoxin A, SEA)-pulsed EBV (Epstein-Barr virus)-transformed lymphoblastoid B cells (B-LCL) expressing high levels of the co-stimulatory molecules CD86 and CD80 or left untreated (Supplementary Fig. 1b-c and Supplementary Fig. 2a-b) . P-CD3ζ (Y142) as a readout of early T cell activation, the highest titrated concentration of SEA (100 ng/mL), and a time point of 45 min was chosen to investigate functional immune synapses (Supplementary Fig. 1 d-e). In total, we screened nine donors in four independent experiments (Supplementary Fig. 2a) and acquired 1,182,782 images (±SEA, Supplementary Fig. 2b). The designed multi-channel panel consisted of brightfield (BF), F-actin (cytoskeleton), MHCII, CD3, and P-CD3ζ (TCR signaling) allowed to capture a wide range of biologically motivated, potentially relevant characteristics of the immunological synapse (Fig. 1a). Dead, deformed, unfocused or cropped cells were removed using a multi-step pipeline (Methods). Additionally, a set of 5,221 images from seven randomly selected donors was labeled by an expert immunologist (K.E.) into nine classes organized in two levels. (Fig. 1b and Supplementary Fig. 2e). The first level represented the number of existing cells in the image: singlets (n=1), doublets (n=2), and multiplets (n>2). The second level characterizes the type of the cells, their interactions to each other and the presence of TCR signaling. The singlets are composed of “single B-LCL”, “single T cell w/o signaling” and “single T cell w/ signaling” classes. The doublets include the “T cell w/ small B-LCL”, “B-LCL and T cell in one layer”, “synapse w/o signaling”, “synapse w/ signaling”, and “no cell-cell interaction” classes. The class “multi-synapse”, contains more than two cells and at least one B-LCL and T cell. Even though the ‘T cell w/ small B-LCL’ and ‘no cell-cell interaction’ classes were artifacts of the experiments, they were annotated to enhance the predictive power of classification models and subsequently filtered out and not used in further analyses (Methods).

**Figure 1:**
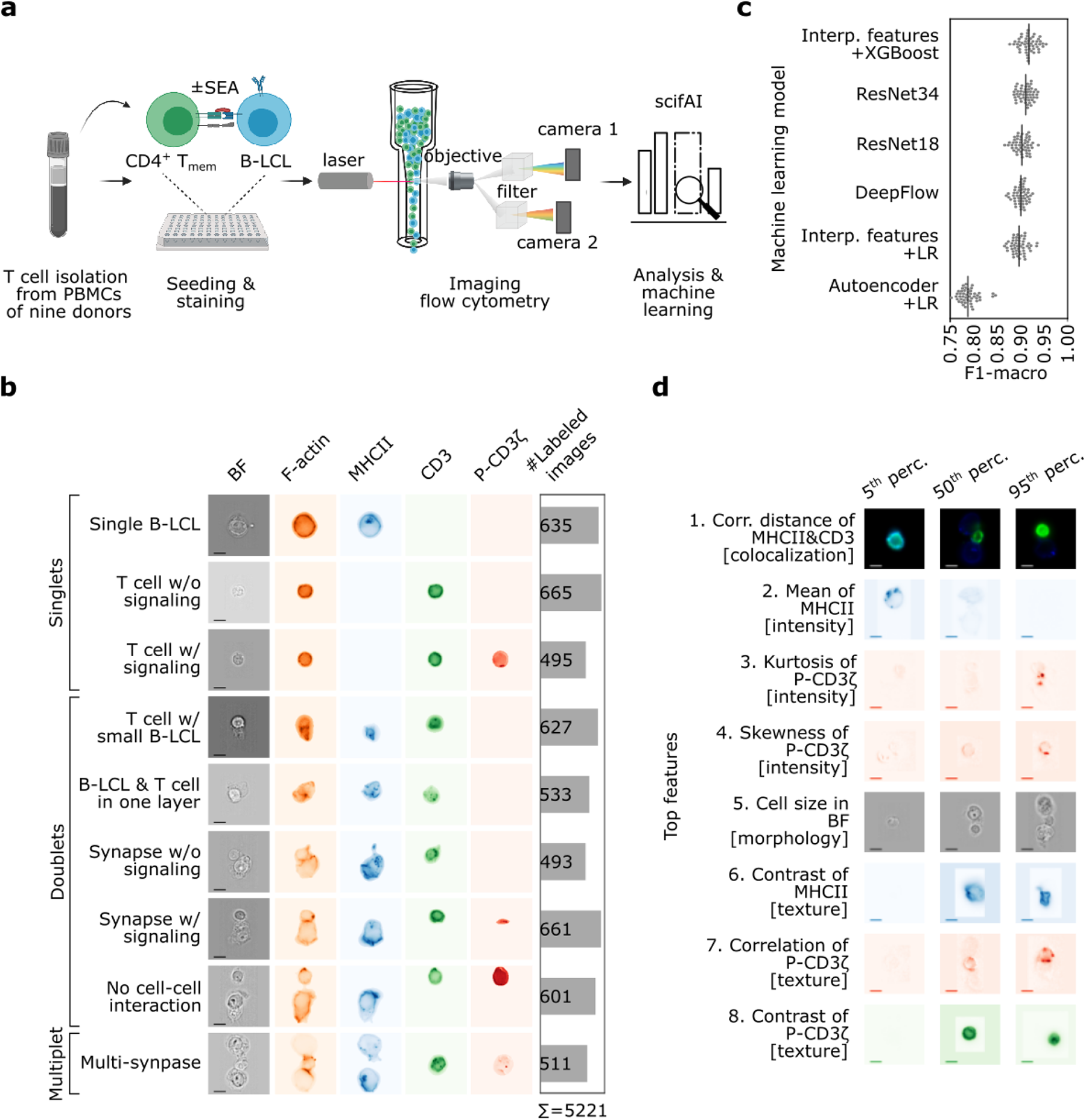
Explainable machine learning accurately predicts immunologically relevant cell classes from IFC data and identifies most informative image features. **a** Schematic representation of the data generation and analysis pipeline. To systematically analyze the immunological synapse of T-B conjugates, 1,182,782 images were acquired with an imaging flow cytometer. Next, scifAI was used to extract morphological features, train machine learning models, profile immunological synapses and characterize the functionality of therapeutic antibodies. **b** A subset of 5221 images was annotated by an expert into nine immunologically relevant classes that can be grouped into singlets (either B or T cells), doublets (with one B and one T cell), and multiplets (containing >2 cells). Cell images show brightfield (BF, scale bar = 2.4μm), F-actin (cytoskeleton), MHCII, CD3 and P-CD3ζ (a marker for TCR signaling). **c** Six different approaches to train predictive machine learning models for the identification of the immunologically relevant classes were benchmarked, combining different classification algorithms and feature engineering strategies. These approaches included interpretable (interp.) features combined with explainable classifiers, an autoencoder to generate data-driven features, an explainable classifier, and three convolutional neural networks. Interpretable features combined with the XGBoost classifier resulted in the best trade-off between interpretability and classification performance. **d** Top eight features for the detection of cell classes were ranked based on in-model feature importance. The features include colocalization, texture and intensity of MHCII, CD3 and P-CD3ζ. The exemplary images are taken from donor 7 in experiment IV (Supplementary Fig. 2) sampling from the 5th, 50th and 95th percentile of the distribution of each feature.

To assess intra- and interrater variability, 100 labeled samples were randomly selected. Three annotators with diverse backgrounds (K.E. – immunology and T cell biology, S.S.B – data analysis, N.K.C. – IFC analysis) annotated the images, yielding Cohen’s kappa ^28^ scores of 0.84 (intra-), 0.79 (inter-), and 0.66 (inter-rater), respectively with respect to the original annotation of the data, which gave us confidence in the reproducibility and validity of the original annotation data.

### scifAI: An explainable AI python framework for the analysis of multi-channel imaging flow cytometry data

As the foundation for all further analysis, we developed the single-cell imaging flow cytometry AI (scifAI) framework. The open-source framework was developed in python, leveraging functionality from state-of-the-art modules, such as scikit-learn, SciPy, NumPy, pandas and PyTorch (Methods), allowing for smooth integration and extension of existing analysis pipelines. Universally applicable for single-cell brightfield or fluorescent imaging projects, the framework provides functionality for import and preprocessing of input data, several feature engineering pipelines including the implementation of a set of biologically motivated features and autoencoder-generated features (Methods), as well as methodology for efficient and meaningful feature selection. Moreover, the framework implements several machine learning and deep learning models for training supervised image classification models, e.g. for the prediction of cell configurations such as the immunological synapse. Following the principle of multiple-instance learning, the framework also implements functionality to regress a set of selected images, against a downstream continuous readout such as cytokine production. Extensive documentation, as well as how to reproduce the analysis in the form of Jupyter notebooks is provided online at https://github.com/marrlab/scifAI/ and https://github.com/marrlab/scifAI-notebooks.

scifAI’s explainable design follows the definition of Singh et al. ^29^ and Tjoa et al. ^30^, where machine learning models are explained by either in-model mechanisms, such as Gini-index, or by using post-model methods, such as saliency maps ^31^ for deep learning models. Using interpretable features with a model whose internal mechanisms can be readily analyzed, scifAI natively provides in-model explainability and is fully compatible with other libraries, such as SHAP ^32^ or Captum ^33^, to provide further post-model explainability.

### scifAI enables high-throughput profiling of the immunological synapse

In order to characterize the immunological synapse in an unbiased fashion, we first designed and computed a series of biologically motivated, interpretable features using the scifAI framework. These features were based on morphology, intensity, co-localization, texture and synaptic features extracted from the 5-panel stained images and their corresponding masks (Methods and Supplementary Fig. 3a-c). Synaptic features were implemented based on the ratio of the signal intensity of each fluorescent channel in the synaptic area to the whole cell.

Leveraging the large amount of unlabeled data, we also implemented a multi-channel convolutional autoencoder to learn a second set of data-driven features from the images in an unsupervised fashion ^24^. The autoencoder was designed to encode the images to a 256-dimensional abstract feature space by reconstructing the input images. Considering the computational complexity of machine learning modeling on high-throughput data, all mentioned algorithms were implemented using parallel computing, fully leveraging high-performance computing (HPC) infrastructure to speed up the calculations by distributing the computational load on multiple CPUs (Methods).

Subsequently, scifAI was used to compose a supervised machine learning pipeline for the classification of the 5,221 annotated images across the nine immunologically relevant cell classes. We trained and benchmarked a series of supervised machine learning models for the prediction of all nine classes using both the interpretable feature space as well as the abstract autoencoder features across all donors and experimental conditions. The models included an XGBoost classifier on the interpretable features and a multi-class logistic regression (LR) on the interpretable and data-driven features. To pre-select the features and reduce the dimensionality, we implemented a feature pre-selection pipeline using an ensemble of different methods (Methods and Supplementary Fig. 4a-b). We also trained a number of convolutional neural network (CNN) architectures such as Resnet18, ResNet34, DeseNet121 and DeepFlow which had previously been shown to be successful in classification tasks on imaging flow cytometry data ^22, 24, 34^. The CNN architectures intrinsically learned a feature representation based on the input images and their corresponding labels. All models were trained on a stratified ±SEA-based subset, comprising 2923 (70%) annotated images. In order to benchmark the classification model and feature space combinations, we compared macro F1 scores on the remainder of images as the ±SEA-based hold-out test set comprising 1567 (30%) annotated images (Methods). The XGBoost model using the interpretable feature set performed best (F1-macro=0.92±0.01, mean ± std 5-fold cross-validation with 10 repetition) among all the classifiers. It was followed by convolutional neural networks ResNet34 (0.91±0.01), ResNet18 (0.90±0.01), and DeepFlow (0.90±0.02). They were followed by multi-class logistic regression using the interpretable feature set (0.90±0.02), and logistic regression using the data-driven feature set (0.79±0.02). Based on the performance and explainability, the XGBoost model was selected as the final classifier for label expansion to the full dataset (Fig. 1c). Investigation of the model’s confusion matrix on the hold-out set revealed that misclassifications occurred mostly within the cell classes’ signaling property, whereas all other classes showed good overall concordance (Supplementary Fig. 4c).

To confirm the validity of the results, a leave-one-donor-out cross-validation using the interpretable features and XGBoost was performed. The cross-validation yielded F1-macro values of 0.88±0.04, demonstrating good generalizability across donors (Supplementary Fig. 4d). After training the XGBoost classifier, we also explored which underlying features drive the class prediction, ranking features by their respective feature importance (Fig. 1d, Methods). The most predictive features were based on the colocalization of CD3 & MHCII, colocalization of MHCII & P-CD3ζ, the texture of MHCII and CD3, and intensity of P-CD3ζ. Based on the features and the definition of classes, one could speculate that the classifier uses (i) the texture of CD3 and MHCII to detect the existence of T and B-LCL cells in the image, (ii) the colocalization of CD3 & MHCII to detect the different doublets types and (iii) the intensity of P-CD3ζ and colocalization of MHCII & P-CD3ζ to detect whether there is a signaling T cell in the image (Fig. 1d).

### A subset of annotated data and available IFC channels suffices for a high classification performance

Considering that manual annotation of images can be time-consuming, we performed an ablation study to investigate how many annotated samples were necessary to achieve a high classification performance. We repeatedly trained the model on stratified subsets of the ±SEA-based training data (5%, 15%, …, 95%) and evaluated the F1-macro on the ±SEA-based test set. The results showed that by using 1500 images (45% of the training data), we could achieve 0.90±0.01 F1-macro on the test set (Supplementary Fig. 5a). This result demonstrated that it is possible to halve the manual annotation time and still achieve a similar quality of classification performance, as compared to using the whole annotated set (0.92±0.01).

Next, we investigated the effects of fluorescent channels on the classification performance to determine which antibodies are essential for the detection of synapses and which ones can be freely exchanged depending on the biological context. Considering that BF is a stain-free channel provided for free by IFC, we kept the BF channel fixed and added all possible combinations of the fluorescent channels to train the model. We found that the combination of the channels BF, MHCII, and P-CD3ζ sufficed to reach an F1-macro of 0.91±0.01 (Supplementary Fig. 5b), similar to using all the channels (0.92±0.01).

### Characterizing the impact of therapeutic antibodies on synapse formation

We next used scifAI to investigate effects of therapeutic antibodies on the formation of the immunological synapse and to better characterize their morphological profiles. This analysis included the investigation of potential class frequency changes and feature differences. We chose two antibodies, one activator and one inhibitor of immune responses. The activating T cell bispecific (TCB) antibody was designed to target CD3 and CD19, a co-receptor of B cells ^35^ (Fig. 2a). The inhibitory antibody, Teplizumab, is described to only bind to CD3 (Fig. 2c) and has been shown to dampen T cell responses ^36, 37^. For each antibody an appropriate control (Ctrl-TCB and isotype) was run within the same experiment and donor. Since Teplizumab required an existing immune response for subsequent inhibition, we used SEA to first stimulate the T cells (Fig. 2c). The same setup was also used for the isotype control. Six donors across two experiments for CD19-TCB and seven donors across three experiments for Teplizumab were measured (Fig. 2b,d and Supplementary Fig. 2c-d). To determine class frequency changes between the antibody and its control, the previous XGBoost classifier (Fig. 1c,d) was used to predict the class for all images based on the interpretable features. A data cleaning pipeline was also implemented to filter out unwanted images such as experimental artifacts (Methods and Supplementary Fig. 6). To ensure that the previously trained XGBoost model was transferable from ±SEA to the antibody experiments, an expert (K.E.) annotated a randomly selected subset of 396 images for CD19-TCB and 227 images for Teplizumab. A high concordance between ground truth annotations and XGBoost predictions on the new experiments (macro F1-score=0.86 for TCB and 0.85 for the Teplizumab) confirmed that the trained model generalizes across experiments and can thus be utilized for further analyses (Supplementary Fig. 7). For a compact representation of class frequency changes, we computed log2-fold changes between the antibodies and their controls. In a second step, we focused on the feature differences of synapses under antibody stimulation and selected all the images predicted as ‘synapses w/ signaling’ for each donor and compared interpretable features from only fluorescent channels including texture, synaptic features, morphology, intensity and co-localization between antibodies and their controls (Methods). Considering that we were interested in the mode of action of antibodies, we focused on fluorescent channels providing targeted information on components of the cell expected to change during synapse formation morphologically.

**Figure 2.**
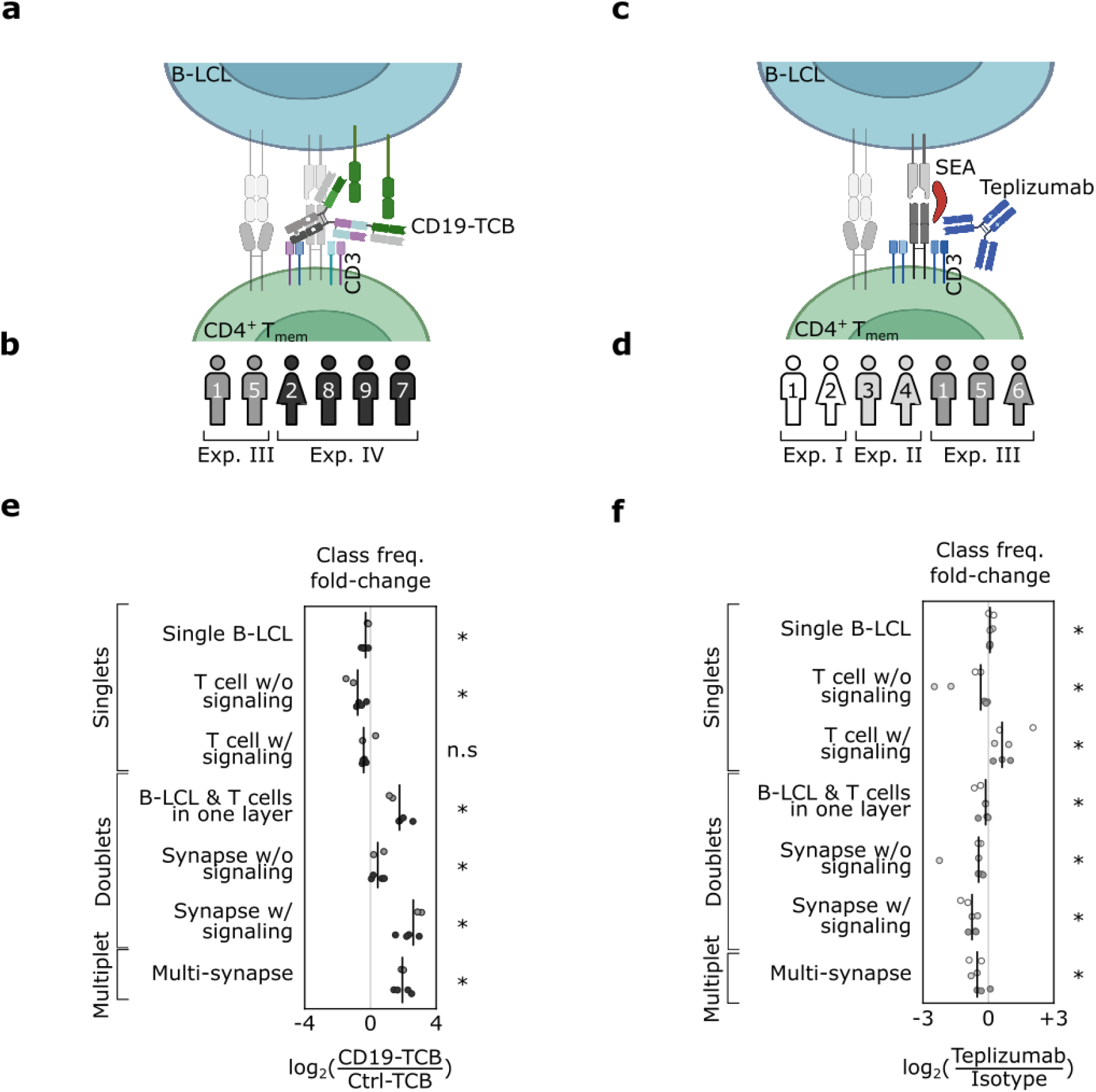
CD19-TCB and Teplizumab show significant changes in the frequencies of synapses. **a, c** Schematic representation of the mode of action of CD19-TCB and Teplizumab. CD19-TCB binds with one arm to the T cell receptor CD3 and with two arms to the B cell co-receptor CD19 thereby bringing T and B cells into proximity and activating T cells. Teplizumab binds with two arms to CD3 and has been described to inhibit T cell activation. To exert their full suppressive function, the T cells needed to be first stimulated via the superantigen SEA. **b, d** Donors and their respective experiments used for the class frequency analysis in Fig 2. e-f, feature difference analysis in Fig. 3, and cytokine prediction analysis in Fig. 4. **e-f** Class frequency differences depicted as log2 fold-changes between CD19-TCB or Teplizumab and their corresponding controls (Ctrl TCB & isotype). Each dot represents a donor color coded as in b or d. The vertical black line is the median across donors for each class.

### CD19-TCB increases the formation of stable immune synapses

Stimulation of the immune response by CD19-TCB led to a significant increase of doublets and multiplets frequencies. The ‘synapse w/ signaling’ class showed thereby the highest increase (median log_2(CD19-TCB/Ctrl-TCB)=2.7, n=6 donors, p=0.036) followed by ‘multi-synapse’ (median=2.03, p=0.036), ‘B-LCL & T cell in one layer’ (median=1.99,p=0.036), and ‘synapse w/o signaling’ class (median=0.59,p=0.036). For the singlets, the overall trend was a decrease in class frequency of ‘single B-LCL’ (median=-0.21, p=0.036) and single T cell w/o signaling (median=-0.77, p=0.036) (Fig. 2e).

Next, we investigated the feature differences in synapses induced by the CD19-TCB (Methods), comparing the 210 interpretable features from all fluorescent channels. We found 210 significantly increased and 163 significantly decreased features out of 210*6=1260 possibilities from combination of features and donors (Fig. 3a). All donors exhibited mostly similar responses towards the stimulation with CD19-TCB. On average 27±4 features were significantly decreased and 33±7 features were significantly increased per donor (dashed lines bottom Fig. 3a). From these features, we were able to find a number of features with similar changes within at least 4 out of 6 donors (Fig. 3a and Supplementary Table 1). We also observed an increase in ‘mean intensity of P-CD3ζ’ with higher enrichment within the synaptic area (Fig. 3b-c and Supplementary Table 1). In addition, we also detected a stronger enrichment of F-actin and MHCII towards the synapse (Fig. 3d-g). Taken together, the observed increase in doublet and multiplet frequencies as well as a stronger enrichment of F-actin and MHCII in the synaptic area indicated an enhanced formation of tight immunological synapses, translating into an efficient TCR signaling. These observations are in line with the mode of action that is already described in general for TCBs, promoting a stable interaction between tumor cells and T cells ^15, 38, 39^.

**Figure 3.**
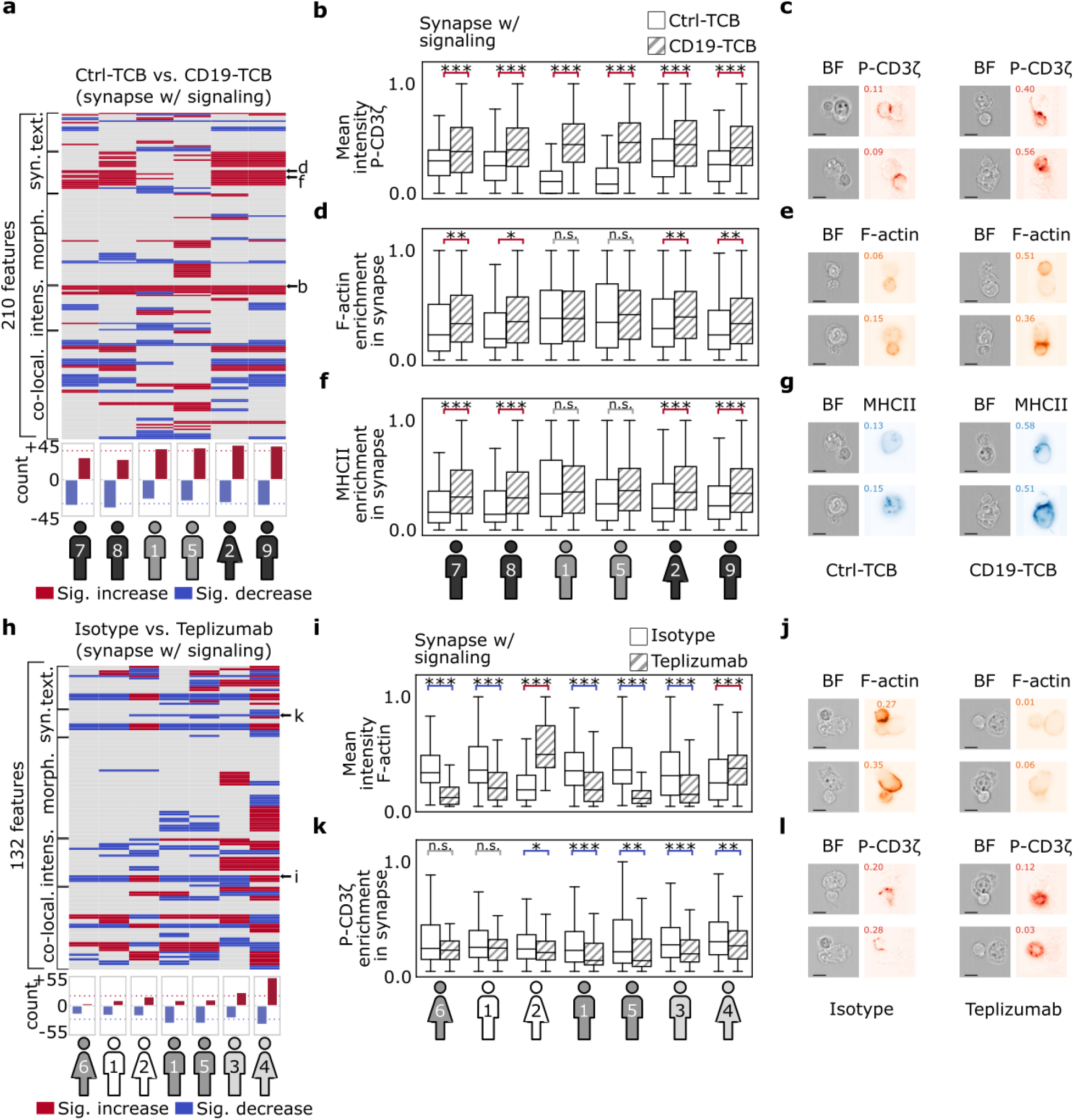
CD19-TCB and Teplizumab induce morphological changes on synapse formation including texture, intensity and synaptic features. **a** Systematic comparison of 210 relevant features between CD19-TCB and Ctrl-TCB across images predicted as ‘synapse w/ signaling’ across six donors. Each line represents a feature and each column represents a donor. For each donor, the features which are significantly increased are depicted with red and the significantly decreased ones are depicted with blue. The donors are sorted based on the number of significantly changed features. The bottom bar plot shows the count of increased or decreased features per donor. Heatmap rows with arrows are shown in detail in **b,d and f**. **b, d, f** Statistical and visual inter-donor comparison of three representative features between CD19-TCB and Ctrl-TCB. For visualization purposes, the features are mapped between zero and one for each donor separately. **c, e, g** Visual representatives for all features were randomly sampled for both Ctrl-TCB and CD19-TCB from donor 9, and were found to be in concordance with the statistical results (scale bar = 2.4μm). **h** Systematic comparison of 132 relevant features between Teplizumab and isotype across images predicted as ‘synapse w/ signaling’ among all six donors. The color code, barplot and sorting are the same as described in **a**. **i, k** Statistical and visual inter-donor comparison of two representative features between Teplizumab and its isotype (also shown by three small arrows in **h**). **j, l** Visual representatives for two features were randomly sampled for both isotype and Teplizumab from donor 3 and were found to be in concordance with the statistical results (scale bar = 2.4μm).

### Teplizumab alters synapse formation and TCR signaling

In contrast to the CD19-TCB, treatment with Teplizumab reduced the frequency of doublets and multiplets significantly (Fig. 2f). The highest decrease was observed for the ‘synapse w/ signaling’ class (median log_2(Teplizumab/Isotype)=-0.75, n=7, p=0.022), followed by ‘multiplets’ (median=-0.69, p=0.036), and ‘synapse w/o signaling’ (median=-0.57, p=0.022). Accordingly, ‘single T cell w/ signaling’ (median=0.70, p=0.022) and ‘single B-LCL’ (median=0.09, p=0.022) were increased significantly as compared to the isotype. Surprisingly, the T cell w/o signaling’ class frequency was significantly decreased (median=-0.30, p=0.022), probably due to the significant increase of ‘single T cell w/ signaling’ (Fig. 2f).

We next investigated feature differences in synapses induced by Teplizumab in seven donors (Methods). From ‘synapses w/ signaling’ images we extracted 132 features based on F-actin, MHCII and P-CD3ζ and their co-localizations. CD3 features could not be included for the analysis because the binding of Teplizumab and the anti-CD3 staining antibody interfere, therefore an anti-CD4 staining antibody was used to identify T cells. We found 131 significantly increased and 169 significantly decreased features out of 132*7=924 possibilities (Fig. 3h and Supplementary Table 2). In particular, Teplizumab, on average, led to 25±8 significantly decreased features and 19±17 significantly increased features per donor (dashed lines bottom Fig. 3h). Donor 6 showed the least number of changes with five significantly increased features. In contrast, donor 4 yielded the highest number of increased features with 50 features, indicating a fair amount of inter donor-variability. We found a set of features which were significantly increased or decreased for at least 5 out 7 donors (Fig. 3i,k). We observed a decrease in the mean intensity of F-actin whereas donor 2 and 4 indicated a significant increase (Fig. 3i,j). This opposite reaction of the two donors could be also detected for other F-actin related features (Supplementary Table 2). Besides the changes in F-actin features, we also detected a significant reduction of P-CD3ζ intensity within the synapse and observed a stronger clustering of TCR signaling around the whole T cell (Fig. 3k,l). In conclusion, we gained new insights into the immunosuppressive mode of action of Teplizumab as we observed a reduction in the number of synapses as well as changes in the F-actin reorganization and P-CD3ζ signaling towards the synapse.

### Morphological profiles of the immunological synapse predict functionality of therapeutic antibodies *in vitro*

Next, the capabilities of scifAI in predicting the functionality of antibodies by analyzing T cell cytokine production were explored. We included an additional antibody in our analysis, the CD20-TCB (Supplementary Fig. 2c and Methods). CD20-TCB is a therapeutic antibody with the same format similar to CD19-TCB but varying target moiety ^15^. CD3 features could not be included for this analysis because the binding of the TCBs interferes with the anti-CD3 staining antibody resulting in erroneously lower CD3 intensity features. Given the richness of interpretable features, we investigated whether it is possible to forecast downstream T cell responses as measured by Granzyme B (GrzmB) for the TCBs. While the IFC measurement was taken after 45 minutes, GrzmB was measured after 24 hours, respectively, using conventional fluorescence-activated cell sorting (FACS) for each donor and condition to address the effects of the antibodies in later time points (Supplementary Fig. 8a). In line with the differences in target expression, the CD20-TCB (20.23±6.32, n=4) showed the highest expression of GrzmB, followed by the CD19-TCB (12.58±3.04) and Ctrl-TCB (1.16±0.51) (Fig. 4a and Supplementary Fig. 8b). A similar pattern was also detected for killing of two tumor cell lines with different expression levels of CD19 and CD20 (Supplementary Fig. 8c-d).

**Figure 4.**
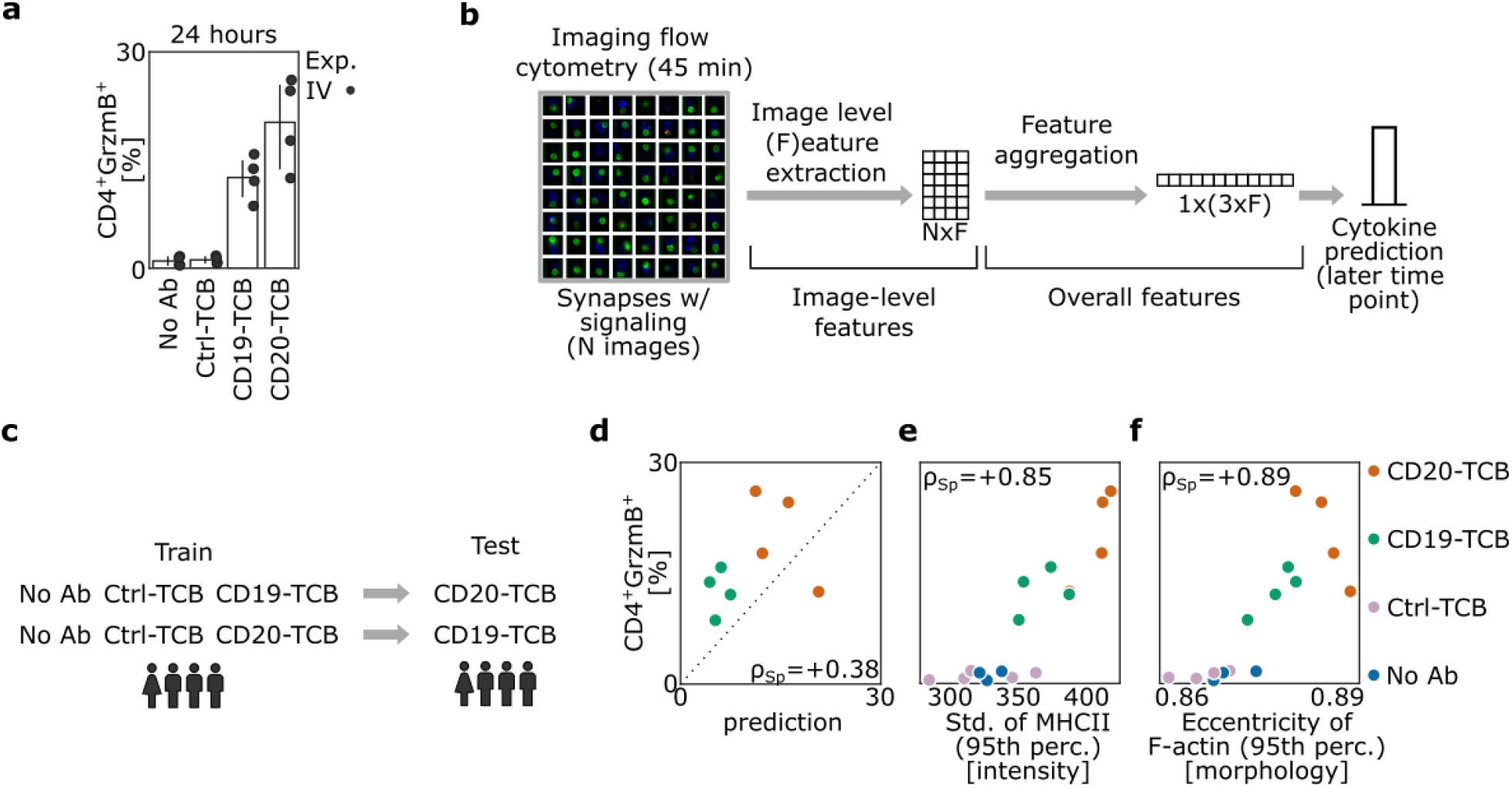
Morphological profiles of synapses are predictive of the functionality of antibodies in vitro. **a** Frequencies of GrzmB^+^ CD4^+^ T cells (gated as single live^+^ cells) measured by FACS after 24 hours. Corresponding to each imaging flow cytometry experiment, a separate FACS experiment was performed with the same batch to obtain the cytokine production values. Each dot represents a donor from experiment IV. **b** Data aggregation pipeline: For each donor and condition, images identified as synapses with and without signaling are selected (N = number of detected images). Then image-level features (F = number of relevant features) are extracted across all selected images. For each donor and condition, the features were aggregated using 5th, 50th and 95th percentile. This 620 dimensional aggregated feature reduces the cytokine prediction to a multivariate regression task. **c** Analysis scheme: Cytokines are predicted for an unseen activator antibody by only using the control (No Ab and Ctrl-TCB) and another activator antibody. Data originates from one experiment with four donors. **d** Scatterplot of the predictions versus ground truth values of GrzmB^+^ CD4^+^ frequencies. The dots are only based on the predictions from (c) therefore only CD19-TCB and CD20-TCB are depicted. **e,f** Scatterplot of the features ‘standard deviation of MHCII (95th percentile)’ and ‘eccentricity of F-actin (95th percentile)’ versus the GrzmB^+^ CD4^+^ frequencies. These two features were selected based on the trained linear model in (d). Both features show high correlation with respect to GrzmB^+^ CD4^+^ frequencies.

Since there is a one-to-many relationship between FACS cytokine measurements and IFC images, where each cytokine measurement corresponds to an IFC cell population consisting of 54,708 images on average, an aggregation pipeline based on the interpretable features was implemented. For each donor and condition, first images predicted as synapses were selected and their previously extracted features were aggregated using the 5th, 50th, and 95th percentile. This aggregation ensured that every feature’s extrema and average expression were captured (Fig. 4b and Methods). Next, it was attempted to predict the cytokines for an unseen antibody. Due to the low number of samples, a linear model with Lasso Lars penalization was used (Methods). The cross validation was performed on CD19-TCB and CD20-TCB, while the No Ab and Ctrl-TCB were kept in the training set (Fig. 4c). The prediction performance reached a Spearman correlation of 0.38 (Fig. 4d). While we could observe subtle differences between the predictions and the ground truth, the model correctly identified the separation between CD20-TCB and CD19-TCB. Furthermore, the model suggested that the ‘standard deviation of MHCII (95th perc.) [intensity]’ and ‘eccentricity of F-actin (95th perc.) [morphology]’ as the most important features (Fig. 4e-f).

## Discussion

In the present work, we established scifAI, a pipeline based on two innovative technologies, imaging flow cytometry, and explainable machine learning to understand the mode of action and predict the functionality of therapeutic antibodies *in vitro*. We analyzed morphological profiles of the immunological synapse to better characterize the mode of action of therapeutic antibodies early after the initiation of an immune response and to apply it to forecast downstream T cell responses. This work is the first coherent functional study using large image data sets and the consequential analytical part to visualize, understand and study synapse formation and its link to predictive features.

We generated the largest publicly available imaging flow cytometry data with over 2.8 million images using human primary immune cells from nine donors in four independent experiments that were treated with various therapeutic antibodies to study synapse formation. The large number of acquired images across multiple experiments provided sufficient statistical power to enable the study synapse formation, a potentially low-probability event, going beyond previous works that did, for example, not consider inter-experiments effects or inter- and intra-donor variability ^21, 40^. While we demonstrate good generalizability of our model, we do not expect that the model trained on our specific IFC data will be predictive without additional training on different datasets obtained with other IFC machines out of the box due to the inherent domain shifts in resolution, magnification, light wavelengths, or focal depth. However, exploiting transfer learning, self-supervised pre-training ^41^, and domain adaptation and generalization techniques ^42^, this dataset will be a valuable resource for future applications, such as transfer to other fluorescent imaging modalities, for example, confocal microscopy, where data can be scarce. Training deep generative models to understand the mode of action of therapeutic antibodies ^43^ is another exciting avenue for future research.

To detect and study immunological synapses, we implemented an interpretable feature extraction and machine learning framework in python by only using well-maintained python modules. scifAI natively implements parallel computing, fully leveraging modern HPC infrastructure and allowing for efficient processing and analysis of high-throughput data by parallelizing tasks such as data preprocessing and model fitting across multiple CPUs. These choices guarantee performance, scalability, and reproducibility and facilitate the deployment into existing workflows, which differs from previous works that use a combination of CellProfiler, R and python for each stage of the analysis ^27, 44, 45^. In this work, we followed the definition of Singh et al. ^29^ and Tjoa et al. ^30^ in the context of the explainability of machine learning and AI. We regard the biologically motivated features as interpretable as they are meaningful and can potentially hint at underlying biological mechanisms. We also considered our XGBoost model explainable as it natively provides a feature importance measure ^32^. This design also enables a deeper understanding of the model’s decision-making using additional methods, such as SHAP ^32^. The insights from the interpretable features and explainable models have the potential to generate new biological hypotheses, which can lead to a better understanding of the underlying mechanisms at play. It is essential to mention that explainable models can offer a helpful intuition only if trained on interpretable features and will fail to provide a meaningful interpretation when trained on abstract features such as the bottleneck layer of an autoencoder. The combination of interpretable features and explainable machine learning enabled us to identify various relevant classes, such as immunological synapses, with state-of-the-art accuracy. It also allowed us to investigate the morphological profiles of the immunological synapse in an unbiased way and characterize the mode of action of antibodies in a biologically relevant context. This methodology is thus a substantial contribution to the field, which has so far primarily focused on performance over interpretability by using ResNet CNN architecture as the backbone ^26, 41^. Notably, the scifAI framework can be easily trained with other fluorescent imaging data. As proof of concept, we provide three examples within the scifAI code repository on how to analyze IFC datasets from Jurkat cells ^34^, white blood cells ^22^, and apoptotic cells ^24^ using scifAI.

To demonstrate the capabilities of the scifAI framework we investigated the effects of two therapeutic antibodies on the immunological synapse, the CD19-TCB and Teplizumab, which are both binding CD3 and have been described to activate and suppress T cell responses, respectively ^35–37^. We found that the CD19-TCB forms more stable immune synapses as indicated by a stronger enrichment of MHCII and F-actin within the synapse that was paralleled by a higher intensity of P-CD3ζ. The formation of stable T cell-tumor cell synapses has already been reported for other TCBs like the CEA- and CD20-TCB ^15, 38^. In contrast to the CD19-TCB, treatment of Teplizumab yielded a decrease in synapse formation and prevented F-actin reorganization as well as localization of P-CD3ζ towards the synapse. These observations gave new insights into the immunosuppressive mode of action of Teplizumab that has rarely been investigated *in vitro* so far ^36, 37^. One could speculate that steric hindrance by Teplizumab prevented T-B cell interactions leading to less stable synapses and reduced cytokine production. It has been shown that antibody size and format can have a substantial impact on synapse formation ^16, 17^. Another hypothesis could be that binding of Teplizumab induced strong TCR internalization that led to diminished SEA-mediated TCR-MHCII crosslinking and thus inhibited T cell activation. The reduced P-CD3ζ intensity in the synaptic area and the observed unpolarized distribution of P-CD3ζ signal around the whole T cell could also indicate altered TCR signaling that might be translated into a reduced T cell effector function. High numbers of peripheral P-CD3ζ microclusters have been already reported for self-reactive T cells with altered synapse formation and aberrant T cell responses ^6^. Interestingly, scifAI identified features within the synapse class revealing inter donor-variability upon stimulation with the different antibodies. However, we were not able to correlate the variability of those features to a different functionality *in vitro* because the differences in the T cell responses between donors were just minor. Patient material from ongoing clinical trials could help to further elaborate if these synapse features could be used to predict clinical response. In that case, scifAI could enable us to rapidly screen for responders *in vitro* and potentially pre-select suitable patients for clinical trials. Taken together, by applying scifAI we were not only able to thoroughly investigate the mode of action of therapeutic antibodies by identifying significant features, but also to gain more insights into inter donor-variability that might potentially translate into different functional outcomes *in vivo*.

The immunological synapse has previously been studied using high-content cell imaging on human cell lines and primary cells with an artificial APC system that utilized plate-bound ICAM-1 and stimulatory antibodies ^45^. Although German et al. convincingly demonstrated the capabilities of their pipeline by profiling the immunological synapse, they did not investigate whether they can use these profiles in predicting drug effectiveness ^45^. In other studies, the potential of synapse formation was also investigated for CAR T cell therapy, where investigators used the mean intensity of stainings such as F-actin and P-CD3ζ per cell, clustering of tumor antigen and polarization of perforin-containing granules as a measure of synapse formation quality. These features varied between different CAR T cells and correlated with their effectiveness *in vitro* and *in vivo* as well as with clinical outcomes ^46, 47^. In our work, we improved this by incorporating 296 biologically motivated features such as texture, intensity statistics and synaptic related features.

This work is the first in using interpretable features of the immunological synapse to predict the effectiveness of therapeutic antibodies on T cell cytokine production. These features allowed us to predict the functional outcome of an unseen antibody and to pinpoint the driving factors required for the prediction. For the TCBs we found intensity of MHCII and morphology of F-actin as the most prominent features in predicting cytokine readouts. These features can be used as inspiration for further investigation and formulation of new hypotheses on the mode of action of TCBs. Nonetheless, additional experiments are necessary to validate them. The ability to predict unseen antibodies could potentially enable the investigation of various antibody formats to better understand mechanistically how different formats can impact T cell responses and help to guide format selection.

scifAI is an end-to-end data acquisition and analysis framework which can be adjusted to investigate various hypotheses and to develop diverse applications based on imaging flow cytometry data. For instance, while in this study memory CD4^+^ T cells were analyzed as they are poised to show faster immune responses and a higher synapse propensity compared to naive T cells ^49^, imaging and analysis of CD8^+^ T cells, as the main players in cytotoxicity, could further elaborate how synapse features correlate with killing efficiency of therapeutic antibodies against tumor cells. scifAI can also be utilized in the design of IFC experiments, optimizing the number and type of stainings, as well as the total number of images per donor to be acquired. In Pharma R&D, scifAI has the great potential to improve the quality and the speed of antibody development, for example giving new insights towards the mode of action of particular candidate molecules, or to predict *in vitro* efficacy in high-throughput. AI assisted identification of lead molecules and better prioritization in terms of epitope, affinity, avidity and antibody format can have a huge impact on the decision making process. Above that, we can foresee that scifAI can even help to identify responders among patient populations and predict their clinical outcomes. In a nutshell, IFAI could provide substantial benefit by assisting the investigation of the mode of action and the functionality of newly generated antibody candidates.

## Acknowledgements

The authors thank Ronan Le Gleut from the Core Facility Statistical Consulting at Helmholtz Munich for the statistical support. We thank Florian Limani from the Roche Innovation Center Zurich for running the killing assays. We acknowledge the Core Facility Flow Cytometry at the Biomedical Center, Ludwig-Maxmilians-Universität München, for providing equipment, services and expertise. We also thank Martin Turner for the fruitful discussions at the beginning of the project that challenged the biological combination of cell markers. Finally, we thank all members of the CD19-TCB and CD20-TCB team as well as colleagues from Large Molecule Research (LMR), especially Alain Tissot, Diana Pippig and Olaf Mundigl, and colleagues from Cancer Immunotherapy, Oncology at Roche Pharmaceutical Research and Early Development (pRED) for their valuable input and scientific discussion.

## Software and data availability

The scifAI code and instructional notebooks on how to run the code and built analysis pipelines can be found under

● https://github.com/marrlab/scifAI/
● https://github.com/marrlab/scifAI-notebooks

The dataset ^48^ can be accessed here: https://doi.org/10.5061/dryad.ht76hdrk7

## Author contributions

KE designed and performed the experiments. SSB wrote the analysis code and created the figures. SSB and KE wrote the manuscript with input from EG, CM and FS. EG, CM and FS conceived the study. NK and FT helped with the manuscript narrative, analysis ideas and editing. SH and MB provided scientific input, designed experiments and supported the writing of the manuscript. All the authors have read and approved the manuscript.

## Materials & Methods

### Cell line culture

EBV-transformed B-lymphoblastoid cell line (B-LCL) from donor 333 was obtained from Astarte Biologics (# 1038-3161JN16) and cells were cultivated in RPMI-1640 medium (PAN-Biotech; cat # P04-17500) with 10% FBS (Anprotec; cat # AC-SM-0014Hi) and 2 mM L-glutamine (PAN-Biotech; cat# P04-80100). Z138 (MCL, gift from University of Leicester) and Nalm-6 (ALL, DSMZ ACC 128) tumor cells were cultivated in RPMI1640 containing 10% FBS and 1% Glutamax (Invitrogen/Gibco # 35050-038).

### Immune synapse formation and imaging flow cytometry

To analyze immune synapses, human memory CD4^+^ T cells were isolated from peripheral blood mononuclear cells (PBMCs) of nine healthy human donors using a negative selection EasySep Enrichment kit from STEMCELL Technologies (cat #19157). Live/dead staining of T and B-LCL cells was separately performed using the fixable viability dye eF780 for 15 min at RT (eBioscience; cat # 65-0865-14). Cells were then re-suspended in RPMI-1640 medium supplemented with 10% FBS (Anprotec; cat # AC-SM-0014Hi), 5% Penicillin-Streptomycin (Gibco; cat # 15140-122) and 2 mM L-glutamine (PAN-Biotech; cat # P04-80100). Afterwards B-LCL cells were transferred into a well of a 96-well round bottom plate (300.000 cells per well) and were pre-incubated with the superantigen Staphylococcal enterotoxin A (SEA) (Sigma-Aldrich; cat # S9399) for 15 min at 37°C or left untreated. Human CD4^+^ T_mem_ were added to the afore-prepared B-LCL cells (250.000 cells per well) to generate a final ratio of 4:3 (B-LCL:T_mem_) and subsequently the appropriate in-house made compounds (10 µg/mL of Isotype Ctrl or Teplizumab and 1 µg/mL (5 nM) of Ctrl-TCB, CD19-TCB^35^ or CD20-TCB^15, 39^) were added to the B-LCL-T_mem_ cell co-culture. To strengthen the conjugate formation between B-LCL and T cells they were centrifuged at 300xg for 30 sec and then directly transferred to a 37°C incubator for 45 min. Thereafter, the medium in each well was carefully aspirated with a pipette and cells were immediately fixed for 12 min at RT followed by permeabilization using the Foxp3/Transcription factor staining buffer set from eBioscience (cat # 00-5523-00). Intracellular staining was performed in permeabilization buffer containing fluorescently-labeled antibodies for 40 min at 4°C: CD3-BV421 (clone UCHT1, Biolegend; cat # 300433), HLA-DR-PE-Cy7 (clone L243, Biolegend; cat # 307616), Phalloidin AF594 (ThermoFisher; cat # A12381) and P-CD3ζ Y142-AF647 (clone K25-407.69, BD cat # 558489). After washing, cells were suspended in FACS buffer (PBS supplemented with 2% FBS) and acquired on an Amnis ImageStream^X^ Mark II Imaging Flow Cytometer (Luminex) equipped with five lasers (405, 488, 561, 592 and 640 nm). On average, around 55,000 images were collected per sample at 60x magnification on a low speed setting. IDEAS software (version 6.2.187.0, EMD Millipore) was used for data analysis and labeling of cells. To identify immune synapses using the IDEAS software the gating strategy in Supplementary Fig. 1a was implemented. Cells were first gated on in-focus live^+^ CD3^+^ MHCII^+^ cells. Within this population images that show single CD3^+^ T cells and single MHCII^+^ B-LCL cells were selected using the area and aspect ratio feature. Next, to exclude non-interacting cells the CD3 intensity within a self-created synapse mask was determined. The synapse mask was defined as a combination of the morphology CD3 and MHCII mask with a dilation of 3. Only synapses that showed a CD3 signal in the mask were gated. Finally, T+B-LCL cells in one layer were excluded by using the height and area feature of the brightfield (BF) and single T-B-LCL synapses were analyzed.

### Conventional flow cytometry

For analysis of cell surface markers, live/dead staining was first performed using the fixable viability dye eF780 for 20 min at 4°C (eBioscience; cat # 65-0865-14). Afterwards, cells were pre-incubated with human Fc-block (BD, cat # 564220) in FACS buffer (PBS supplemented with 2% FBS and 1 mM EDTA) for 10 min at 4°C and then stained with the appropriate fluorescently-labeled antibodies for 30 min at 4°C: CD4-BV510 (clone RPA-T4, Biolegend: cat # 300546) or CD4-BV421 (clone RPA-T4, BD; cat # 562424), HLA-DR-PE-Cy7 (clone L243, Biolegend; cat # 307616), CD69-PE (clone FN50, Biolegend; cat # 310906), CD80-APC (clone 2D10, Biolegend cat # 305220 or CD86-PE (clone IT2.2, Biolegend cat # 305406). For intracellular cytokine staining cells were first treated with GolgiPlug (BD Biosciences; cat # 555029) and GolgiStop (BD Biosciences; cat #554724) for at least 2-4 h before being stained. After incubation live/dead staining was performed using the fixable viability dye eF780 for 20 min at 4°C (eBioscience; cat # 65-0865-14). Cells were then fixed and permeabilized using the Foxp3/Transcription factor staining buffer set from eBioscience (cat # 00-5523-00) as described for the synapse formation assay. Intracellular staining was performed in permeabilization buffer containing the fluorescently-labeled antibody Granzyme B-PE-Cy7 (clone QA16A02, Biolegend; cat # 372214) or TNF-ɑ (clone MAb11, BD; cat # 554514) for 30 min at 4°C. Finally, cells were suspended in FACS buffer (PBS supplemented with 2% FBS and 1 mM EDTA) and acquired on a FACS Celesta from BD Biosciences.

A representative example of the gating strategy used for analyzing conventional flow cytometry data in this study is shown in Supplementary Fig. 9. Briefly, lymphocytes were selected in the FSC-A and SSC-A gate. In the next step, single cells were selected using FSC-H/FSC-W and viable cells were identified using the fixable viability dye eF780 (gated on eF780 negative cells). Finally, cells were gated on CD4^+^ T cells and markers of interest were analyzed (see Supplementary Fig. 1d and Supplementary Fig. 8a).

### Tumor Cell Lysis Assays (*in vitro*)

B cell-depleted PBMCs derived from blood of healthy donors were prepared using standard density-gradient isolation followed by B cell depletion with CD20 Microbeads (Miltenyi; cat # 130-091-104). B cell-depleted PBMCs were then incubated with the tumor targets (Z-138 or Nalm-6) at a ratio of 5:1 for 24 h in the presence or absence of CD20-TCB or CD19-TCB. Tumor cell lysis was calculated based on LDH release (LDH Cytotoxicity Detection Kit from Roche Applied Science) and normalized to spontaneous release (PBMCs + targets without treatment = 0 % tumor cell lysis) and maximal release (lysis of tumor targets with Triton X-100 = 100 % lysis).

### Quantification of CD20 and CD19 expression

CD19 and CD20 expression on B-LCL cells were determined using the Quantum™ Alexa Fluor® 647 MESF Kit from Bangs Laboratories (cat # 647) according to the manufacturer’s instructions using an anti-human CD20-AF647 (Biolegend # 302318) or and anti-human CD19-AF647 (Biolegend # 302220) antibody as well as the corresponding isotype controls (muIgG1 (Biolegend # 400130) and muIgG2b (Biolegend # 400330). For the quantification of CD19 and CD20 molecules on the tumor target cell lines Nalm-6 and Z-138 the QiFi Kit from Dako (cat # K0078) was performed according to the manufacturer’s instructions by using an anti-human CD20 purified (BD # 555621) or and anti-human CD19 purified (BD # 555410) antibody as well as the corresponding isotype controls (muIgG1 (BD # 554121) and muIgG2b (BD # 557351).

### Preparation of the imaging dataset for analysis

We recorded 2,899,575 images from the commercial imaging flow cytometer, Luminex Amnis ImageStreamX Mark II Imaging Flow Cytometer, with estimated throughput of 100-200 events/sec. The dataset consists of nine distinct donors across four independent experiments. Donor 1 and Donor 2 were used twice (Supplementary Fig. 2a). Different conditions were measured which included -SEA (total images=625,001), +SEA (557,781), Ctrl-TCB (330,000), CD19-TCB (324,020), CD20-TCB (254,398), Isotype (405,000), and Teplizumab (403,375). The images contained brightfield (BF), F-actin, MHCII, CD3, P-CD3ζ, and Live/Dead stainings. The Live/Dead staining is only used to filter out the dead cells. For each experiment, the images were compensated using a compensation matrix derived from stained single cells. After the compensation, the raw images (16-bit) and their corresponding channel-wise segmentation masks were exported from the IDEAS software and saved in an HDF5 format. To enable parallelization, each image and its corresponding mask were saved separately. In addition, our expert annotated a subset of data for -SEA (labeled images=1160), +SEA (4061), CD19-TCB (396) and Teplizumab (227). The combination of the ±SEA-based was used for training the classification models where 70% was used for training and 30% for test.

#### Interpretable feature engineering from images

We extracted a set of 296 biologically motivated features to study the immunological synapse. These features included morphology, intensity, co-localization, texture and synaptic related values (see Supplementary Fig. 3). The morphology features were calculated based on the segmentation mask from each channel. The features included ‘area’, ‘bounding box area’, ‘convex area’, ‘eccentricity’, ‘equivalent diameter’, ‘Euler number’, ‘extent’, ‘maximum Feret diameter’, ‘minimum Feret diameter’, ‘filled area’, ‘length of major axis’, ‘length of minor axis’, ‘Hu moments’, ‘orientation’, ‘perimeter’, ‘Crofton perimeter’, ‘solidity’, ‘weighted Hu moments’. All the morphology features are extracted using scikit-image library ^49^. For the intensity features, first the cells were segmented using their corresponding mask. The intensity features included ‘min’, ‘sum’, ‘mean’, ‘standard deviation’, ‘skewness’, ‘kurtosis’, ‘max’ and ‘Shanon entropy’. In addition, the percentile of intensity values including ‘10th percentile’, ‘20th percentile’, …, ‘90th percentile’ were calculated. All of the intensity features were calculated based on NumPy ^50^ and SciPy ^51^ functionality. For co-localization features, we implemented ‘dice distance’ and ‘Jaccard distance’ to calculate the masks overlap between two channels using the SciPy ^51^ library. In addition, we calculated the ‘correlation distance’ ^51^, ‘Euclidean distance’ ^51^, ‘Manders overlap coefficient’ ^52^, ‘intensity correlation quotient’ ^52^, ‘structural similaity’ ^49^ and ‘Hausdorff distance’ ^49^. For texture features, we used Gray Level Co-occurrence Matrix (GLCM) features ^53^ including ‘contrast’, ‘dissimilarity’, ‘homogeneity’, ‘ASM’, ‘energy’ and ‘correlation’. The synapse related features were defined as ‘enrichment of Ch (mean)*mean (intensity of Ch in synapse)/mean(intensity of Ch)* of Ch (sum)’= *sum (intensity of Chh in synapse)/sum(intensity of Ch)*, and ‘enrichment of Ch (max)’ = *max(intensity of Ch in synapse)/mean(intensity of Ch)*^40^. Finally, we implemented ‘background mean’ and ‘gradient RMS’ for quality control of images. All these features were implemented using NumPy(version=.18.5), Pandas (1.1.5), SciPy (1.8.0), scikit-image (0.19.2), and scikit-learn(1.0.2)^54^.

#### Autoencoder feature extraction

To leverage the large amount of unlabeled data, we implemented and trained a multichannel autoencoder ^24^. This autoencoder included a separate encoder for each channel. The encoders were designed to map each channel to a 32 dimensional vector. The concatenation of these vectores led to a 5*32 dimensional space. Then these features were mapped to a 256 dimensional feature vector. A decoder on top of the concatenated vectors was implemented for reconstructing the original image. Mean squared error (MSE) was used as the reconstruction loss. The augmentations used for training the autoencoder included random rotation, random scaling, random flipping, random gaussian noise.

#### Feature pre-selection

Considering that the number of features was large, we implemented a feature pre-selection pipeline to select the most relevant features. We followed the work of Haq et al. ^55^ (see Supplementary Fig. 4a). First, the Pearson correlation between the features was measured. If at least two features were highly correlated (|corr|>0.95), then only one of them was kept (at random), and the rest were eliminated. In the next step, six different methods were used to rank the features. These methods included mutual information, linear support vector machine, logistic regression with L1 regularization, logistic regression with L2 regularization, random forest, and XGBoost. The *top-k* (hyper-parameter to be selected) features from each method were selected, and their union was used. After this reduction, the Spearman correlation matrix between the features was calculated, and spectral clustering was performed on the correlations. Then, *m* clusters were created, and one feature at random per cluster was selected. The last step was performed to account for multicollinearity between the features.

### Classification

There are three main approaches which are used for training a supervised learning algorithm in this work, feature based approach and deep learning.

#### Classical supervised learning models

We used two different algorithms for training machine learning models. We used a boosting method called XGBoost ^56^ which uses an ensemble of trees on the data (n_trees=100). The second model was a logistic regression. The advantage of using these models was that they provide explainability after the training.

#### Convolutional neural networks

For training supervised deep learning models, we used well-known architectures in the field of computer vision including ResNet18, Resnet34, ResNet50, ResNet152, DeseNet121 and DeepFlow ^22, 24, 34^. All models were pre-trained on ImageNet. Considering these models are originally designed for three RGB channels input, we had to substitute the first convolutional layer with three input channels to five input channels including BF, F-actin, MHCII, CD3 and P-CD3ζ. In addition, the classification layer also needed to be adjusted to have nine classes. However, the rest of the networks were untouched with their predefined ImageNet weights. We used multi-class cross entropy loss for training. The learning rate (lr) was set to 0.001, with adaptive strategy of reducing on plateau of 10 epochs. The augmentations used for training included random rotation, random scaling, random flipping, random gaussian noise. 15% of the training data was selected randomly and used as a development set to avoid overfitting.

#### Classification feature importance

As mentioned, a feature pre-selection filtering was used to reduce the number of features and then an XGBoost classifier was trained on the annotated data. While the XGBoost can provide in-model feature importance, these importances can be biased due to different reasons such as correlation between the pre-selected features, number of features, the pre-selection process, outliers, etc. To account for this, we splitted the training data randomly to 5-folds (stratified) and trained the XGBoost classifier five times, each time using 4 out of the 5 folds. We repeated this process 100 times, leading to 500 different models. In each training, we used a random number of pre-selected features (top-k) between 30 to 200 features. Eventually, for every feature a series of feature importances were obtained. The median feature importance for each feature was used to rank the features.

### Classification performance validation and generalizability

We validated the prediction performance of classification models across multiple datasets. To offer a comprehensive view of the generalizability of our results, we present a complete list of all the steps taken. The first validation was performed on the ±SEA-based test set (30% hold-out set) to confirm inter-experiment comparability (Supplementary Fig. 4c). The second validation was performed using leave-one-donor-out cross-validation to confirm inter-donor generalizability (Supplementary Fig. 4d). Next, the model was validated on two separate test sets with Teplizumab-based and CD19-TCB-based perturbations to validate inter-experiment and inter-stimulation generalizability (Supplementary Fig. 7). Overall, we demonstrated that the XGBoost model trained on ±SEA perturbations could indeed generalize to new perturbations with previously unseen antibodies and inter-donor variability.

### Classification staining importance

To determine which staining contributes the most to the predictions, we used recursive channel elimination. In every run, we always kept BF as it is stain-free. Then train the ‘interpretable features + XGBoost’ based on the features of the selected channels (see Supplementary Fig. 5b).

### Data cleaning pipeline

For each donor, we used the trained XGBoost classifier to predict the class of every image. We excluded low quality and outlier images based on the following data cleaning protocol, where we first describe the types of cells we filtered out, followed by the concrete feature based rule:

1. Dead cells (using Live/Dead staining): ‘mean Live/Dead intensity’ >= ‘mean Live/Dead intensity (90th perc.)’
2. Out-of-focused images: ‘Gradient RMS BF’ > ‘Gradient RMS BF (2nd perc.)’ & ‘Gradient RMS BF’ < ‘Gradient RMS BF (90th perc.)’
3. High entropy images, based on the XGBoost predictions: entropy > 1.0. The entropy was calculated using the SciPy package. This step is done to omit images that the classifier is the most uncertain in terms of prediction.
4. Images predicted as ‘B-LCL’ with low MHCII: ‘mean intensity of MHCII’ < ‘mean intensity of MHCII (5th perc.)’. This step guarantees that the images predicted as ‘B-LCL’ contain a minimum of MHCII intensity.
5. ‘B-LCL’ with ‘area of MHCII’ < ‘area of MHCII (10th perc.)’. This step guarantees that the images predicted ‘B-LCL’ contain a cell with appropriate size.
6. ‘T cell’ with ‘mean intensity of CD3’ < ‘mean intensity of CD3 (1st perc.)’. This step guarantees that images predicted as ‘T cell’ contain a minimum CD3 intensity.
7. ‘B-LCL and T cell in one layer’ with small ‘B-LCL’s: ‘B-LCL and T cell in one layer’ with ‘area of MHCII’ < ‘area of MHCII (20th perc.)’. This step is performed to omit missclassified ‘B-LCL and T cell in one layer’ with small ‘B-LCL’s.
8. Outlier detected via isolation forest ^57^: We used n_estimators=100, max_samples=’auto’, contamination=’auto’, and max_features=20 as the main parameters. For reducing the run time, we only used top 30 features based on feature importance from the XGBoost training.
9. Outlier detected via DBSCAN ^58^: First we transformed all images to 2D dimensional space using Uniform Manifold Approximation and Projection (UMAP). The features were standardized using the mean and std of each feature. For reducing the run time, we only used top 30 features based on feature importance from the XGBoost training. Then a DBSCAN algorithm was run with eps=0.09 and min_samples=5. The resulted clustered were filtered out if (#images in cluster)/(#total images) < 0.0001.

All these steps are based on scikit-learn implementations. All parameters were set using the default value of scikit-learn unless stated otherwise. 20 random examples of filtered out images are shown in Supplementary Fig. 6.

### Class frequency analysis

After the data cleaning, the frequency of each class was calculated with ‘F_C=(#images predicted as C)/(#total images)’ for each condition per donor. To deal with the compositional nature of the data, we used log_2(F_C_antibody/F_C_control) to compare the frequency fold-changes. . This transformation has the advantage that the frequencies do not sum to a constant value. After calculating the log_2 fold-changes, we used the Wilcoxon-rank-sum test for analyzing the effects of antibodies on class frequencies. Wilcoxon-rank-sum tests whether two samples are likely to derive from the same population. To account for multiple testing, we used Benjamini-Hochberg correction for +SEA/-SEA, CD19-TCB/Control TCB and Teplizumab/isotype respectively. Because the experiments were performed independently, we only corrected each of these comparisons separately.

### Feature difference analysis

We analyzed the effect of perturbation with CD19-TCB, and Teplizumab on signaling synapses using the previously trained XGBoost model from the ±SEA training set and then used statistical tests to investigate the mode of action of these therapeutic antibodies. First, we selected the images predicted as ‘synapse w/ signaling.’ We only focused on fluorescent channels as they contain targeted information on components of the cell expected to morphologically change during synapse formation. This procedure yielded 210 features for comparison for TCB based on F-actin, MHCII, CD3, and P-CD3ζ. For Teplizumab, CD3 was not available for the analysis because of the usage of CD4 in recording images for Teplizumab instead of CD3. Thus we analyzed 132 features extracted from F-actin, MHCII, and P-CD3ζ. After the feature selection, we compared the features using the Mann-Whitney U test for each condition and its control. To understand the direction of change, we used the difference in the median of features for each condition and its control. To account for multiple testing, we used the Benjamini-Hochberg procedure with α=0.05. As the conditions were independent, we corrected the p-values for each condition and its control separately. If the test was not significant, we assigned that feature 0 (gray in Fig. 3a,h and 0 in Supplementary Tables 1-2). If it was significant and the median value of the feature for the antibody was greater than the median of the feature of the control antibody, we assigned +1 (red in Fig. 3a,h and +1 in Supplementary Tables 1-2). On the contrary, if the test was significant, but the median value of the feature for the antibody was smaller than the median value of the feature of the control, we assigned -1 (blue in Fig. 3a,h and -1 in Supplementary Tables 1-2).

### Validation of the biological findings

In the study of the mode of action of antibodies, six donors for the CD19-TCB and seven donors for Teplizumab were analyzed, respectively. To avoid systematic bias in the analysis due to potential batch effects and emphasize the generalizability of the results, we included at least two independent experiments for each antibody: experiments III and IV for CD19-TCB and I, II, and III for Teplizumab. This allowed us to focus on consistent findings between the experiments. All statistical tests were corrected for multiple testing to account for the inflated occurrence of false positive results. Finally, visual inspections of the acquired images were done by expert immunologists, and the findings were validated in the literature.

### GrzmB prediction and feature ranking

To predict GrzmB, we only used images predicted as ‘synapse w/o signaling’ and ‘synapse w/ signaling’ for each condition. This choice was done as it is assumed that the synapses will lead to cytokine production. Considering that for each donor and condition we had thousands of images, we used an aggregation pipeline to create a feature vector corresponding to each donor and condition. To reduce the number of features, we only used the consistent feature changes for the CD19-TCB (Fig. 3). For each donor and condition, we aggregated the features using 5th, 50th and 95th percentile to capture the extremes and average of every feature.

After deriving the aggregated features, we used a one-donor-leave-out cross validation to train a linear regression model with LassoLars. The most important features were based on the magnitude of the coefficients.

### Visualizations and Tables

For visualizing the conjugates and biological context, we used www.BioRender.com. For the plots and images, we used matplotlib (version=3.3.2) and seaborn (0.11.2) in Python. Finally, Inkscape was used for creating the figures and tables, including the glossary of all the abbreviations in Supplementary Table 3.

### Reproducing the results

For reproducing the results or running exemplary code provided as part of the software package, it is necessary to download the dataset and install scifAI. Extensive documentation has been provided that will allow users with basic programming knowledge to follow four application examples and, with minor modifications, adapt the code to their needs. To tackle more advanced use cases, such as defining a new feature, or applying scifAI to a completely new data set, more programming experience is required.

---

**Supplementary Figure 1:**
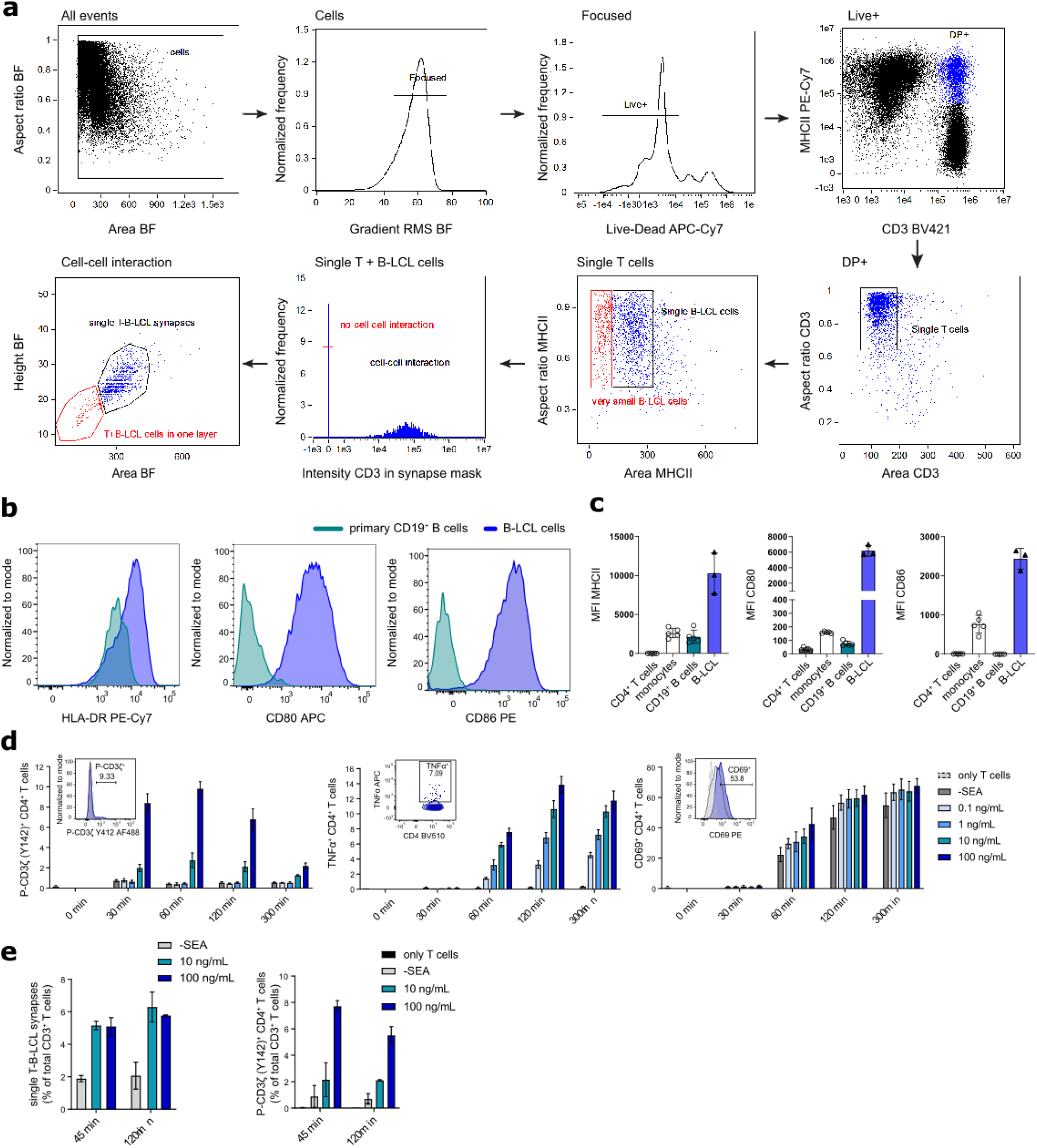
Assay conditions to analyze immune responses between primary human CD4^+^ T cells and B-LCL cells using conventional and imaging flow cytometry. **a** Gating strategy to identify single interacting T-B-LCL synapses using the IDEAS software of the imaging flow cytometer. **b** FACS histograms showing the surface-expression levels of MHCII (HLA-DR), CD80 and CD86 of primary CD19^+^ B cells from PBMCs and B-LCL cells. **c** Quantification (MFI) of MHCII, CD80 and CD86 surface expression on primary CD4^+^ T cells, CD19^+^ B cells and monocytes from PBMCs and B-LCL cells. Data are from five donors in three independent experiments. The B-LCL cell line was analyzed in triplicates. **d** Pre-testing of assay conditions using conventional FACS. Primary memory CD4^+^ T cells isolated from PBMCs of healthy donors were stimulated with B-LCL cells in the presence of different concentrations of SEA (0.1-100 ng/mL) or left untreated (-SEA). Frequencies of P-CD3ζ^+^, TNF-ɑ^+^ and CD69^+^ CD4^+^ T cells were determined at various time points. The small FACS histograms in the bar graphs show the expression levels of the three markers by comparing the highest concentration of SEA (100 ng/mL) with the untreated control (-SEA) after 60 min. The data shown represents one experiment using T cells from three different donors. **e** Percentage of single T-B-LCL synapses and P-CD3ζ^+^ CD4^+^ T cells measured by imaging flow cytometry between two different SEA concentrations (10 and 100 ng/mL) after 45 and 120 min. Data represents two donors.

**Supplementary Figure 2.**
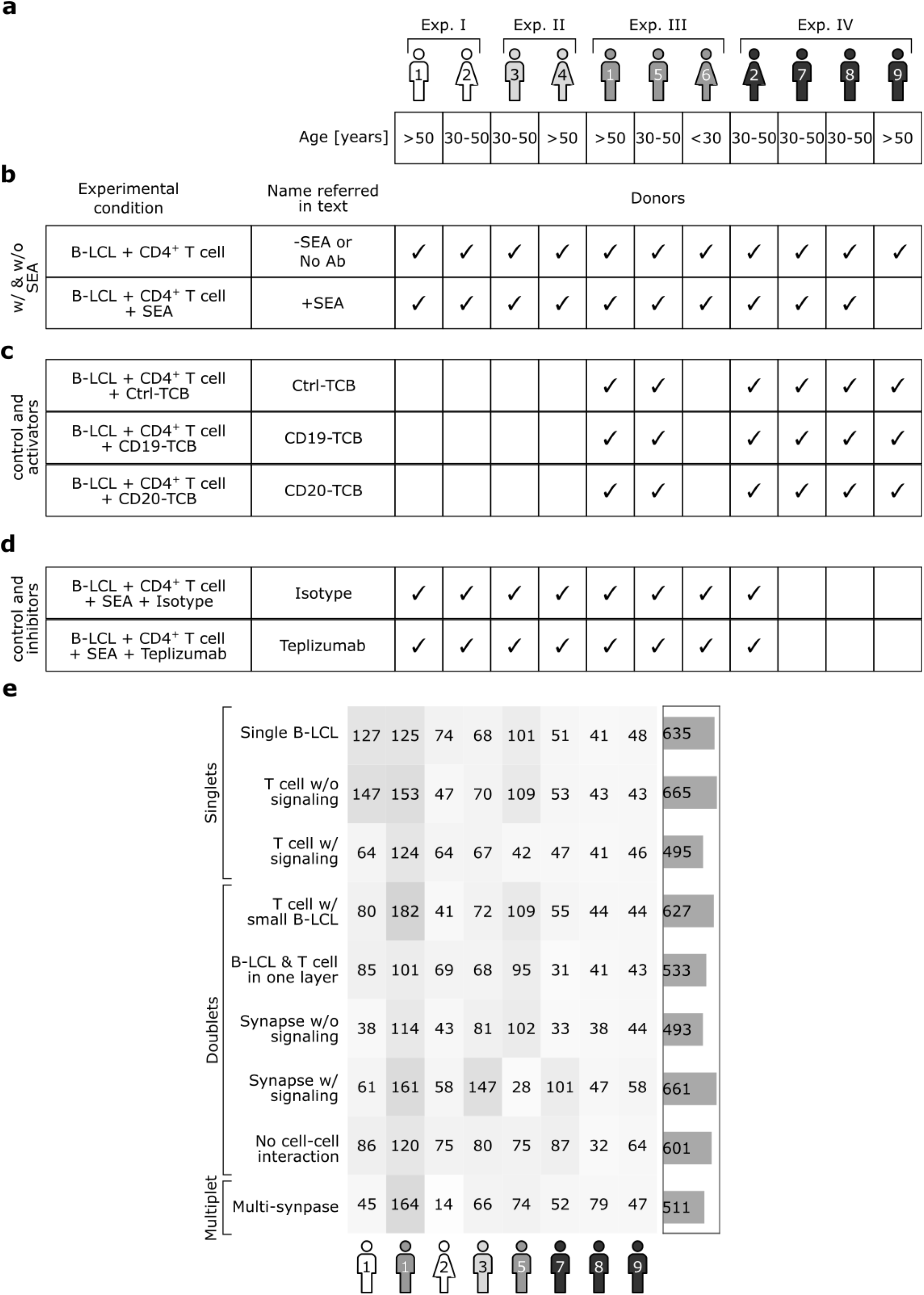
Donors and experiments information. **a** List of the donors, their age, gender and experiment numbers which are used in this study. **b** List of experiments and donors with and without SEA. **c** List of experiments and donors with TCBs and their control. **d** List of experiments and donors with Teplizumab and isotype control **e** Number of labeled by expert data per donor.

**Supplementary Figure 3.**
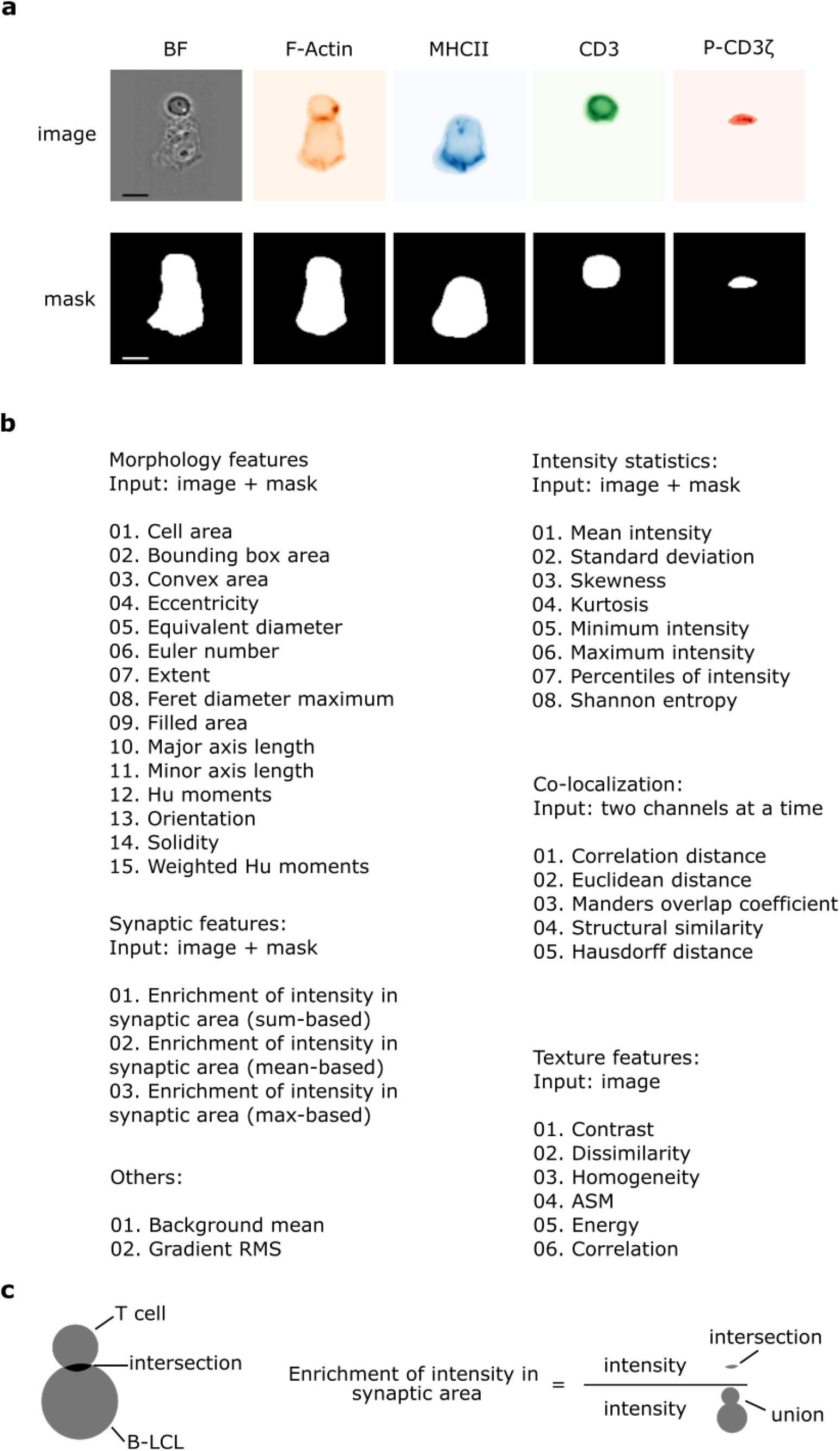
scifAI interpretable feature list. **a** Visual representation of each multi-channel image and corresponding masks. Masks were exported along with images from the IDEAS software (scale bar = 2.4μm). **b** List of all features implemented in scifAI. The morphology, intensity statistics, textures and synaptic features are based on one channel. The colocalization features are based on two channels. scifAI automatically detects the existing channels and generates the specified features. **c** Visual representation of how synaptic features are calculated. To calculate the enrichment of intensity in the synaptic area, the ratio of intersection over the union of cells was used. The intersection is calculated using the masks from CD3 and MHCII channels. For the sum-based enrichment feature, the sum of intensities in each region is calculated. The same logic applied to the mean-based and max-based enrichment features.

**Supplementary Figure 4.**
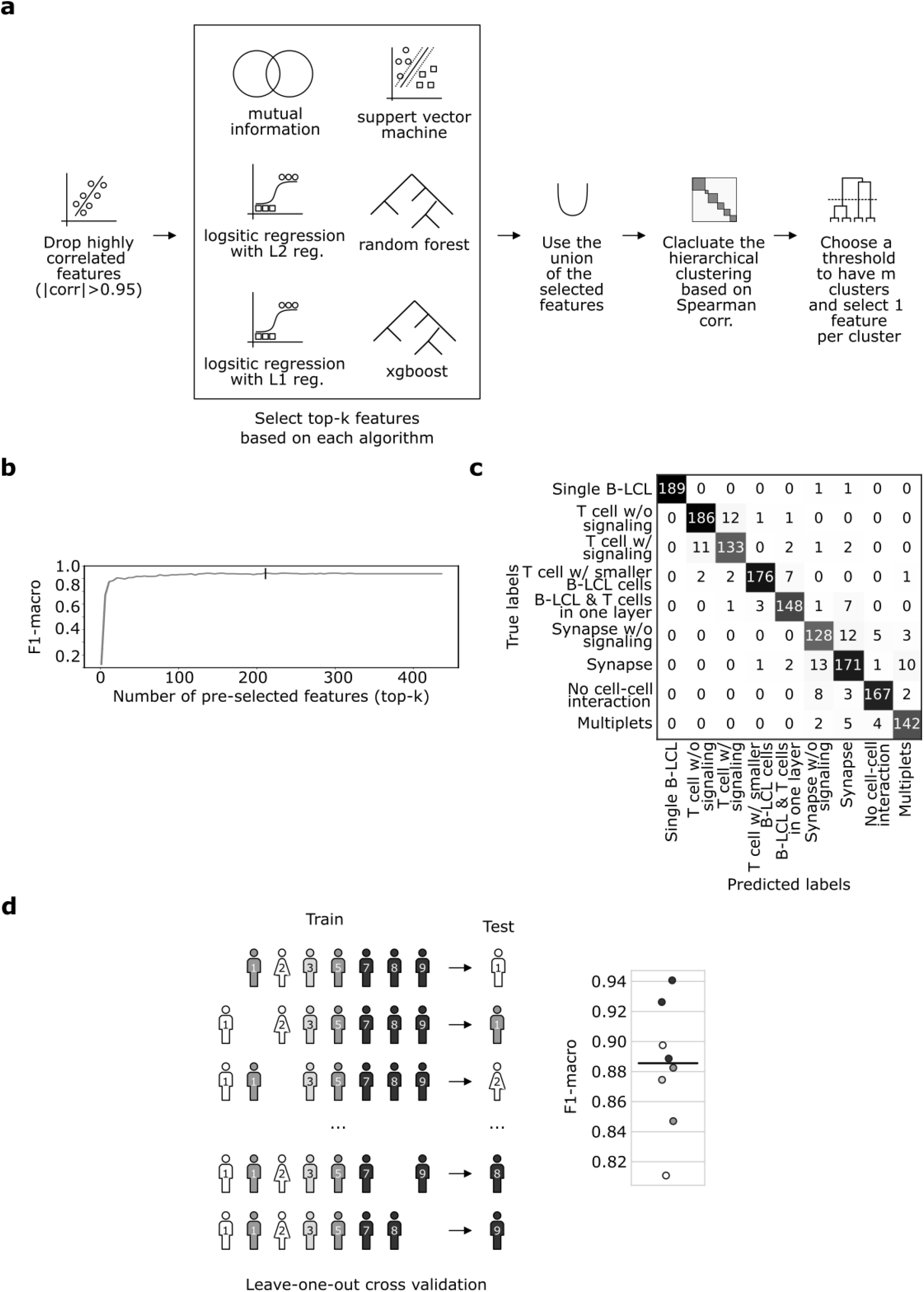
Machine learning pipeline for classification. **a** Feature pre-selection pipeline to reduce the dimensionality of the feature space and removing multicollinearity. First the highly correlated features are dropped. Then an ensemble of different classifiers is trained on the data, and their top-k features are selected. Finally, hierchical clustering is done on top of the union of the features to account for multicollinearity. **b** The optimal number of pre-selected features before passing the pre-selected features to the XGBoost classifier. For obtaining the optimal top-k the data selection pipeline + XGboost was trained on stratified randomly selected 85% of the training set and tested on the rest 15%. **c** Confusion matrix of the data selection pipeline (top-k = 211) + XGBoost, based on the predictions on the test set. **d** Leave-one-donor-out cross validation confirms that the result in c is robust (compared to stratified ±SEA-based train and test set)

**Supplementary Figure 5.**
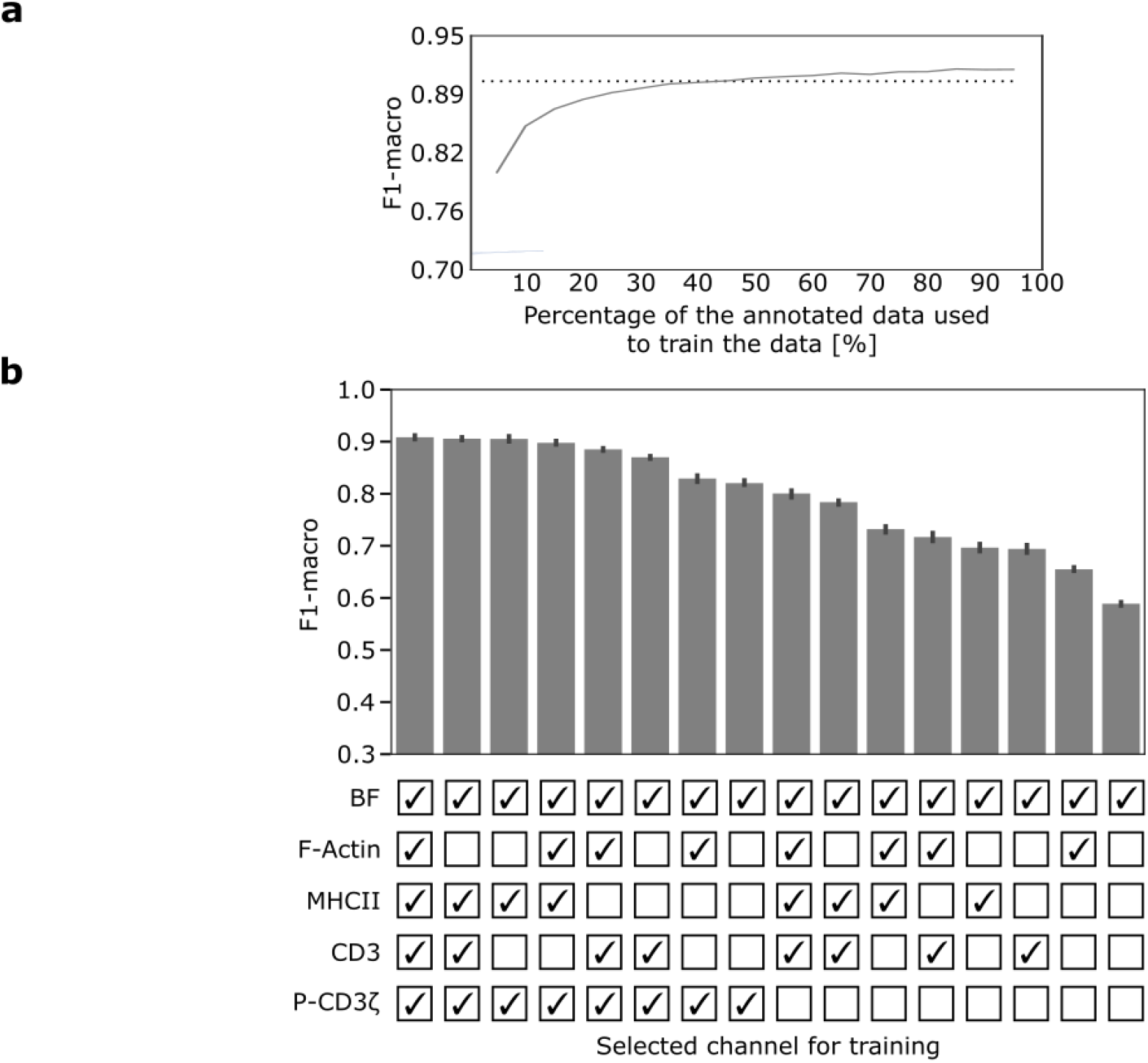
Extracted information in the annotated data. **a** Number of annotated images vs. the classification performance **b** The same XGBoost model was used to train the classifier. In each step, features based on the selected channels were used for training the classifier. As brightfield (BF) is a stain-free channel, it is always kept in the data. The combinations are ranked based on F1-macro. The combination of BF, MHCII and P-CD3ζ (third from left) reaches a similar performance in comparison to using all the channels.

**Supplementary Figure 6.**
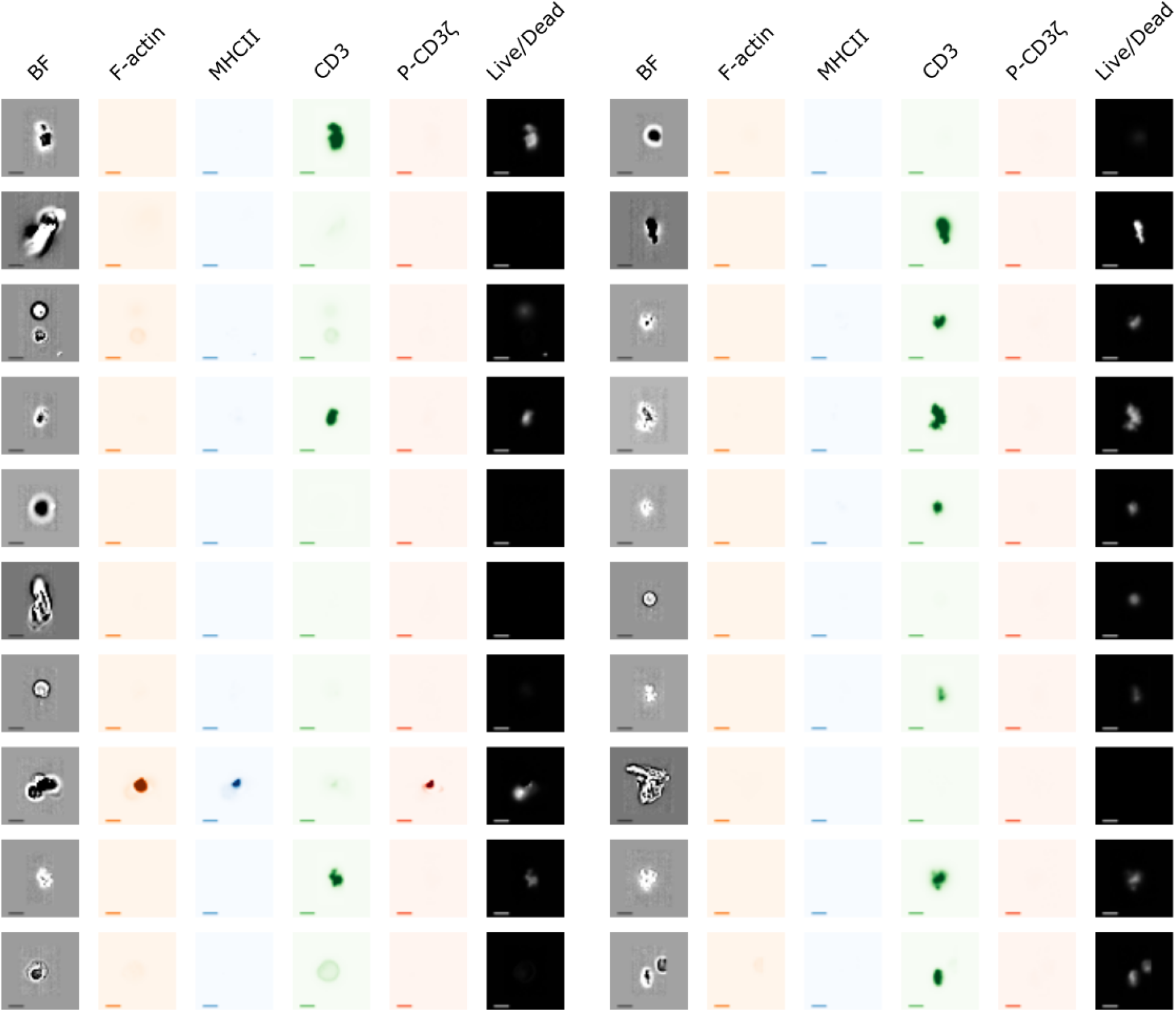
Filtered-out images. 20 randomly selected examples that were excluded from the analysis based on the data cleaning pipeline described in Methods. The examples show dead cells (with signal in the “Live/Dead” channel), out-of-focus images and images with irregular shapes in the brightfield channel, or missing fluorescent expression (scale bar = 2.4μm)

**Supplementary Figure 7.**
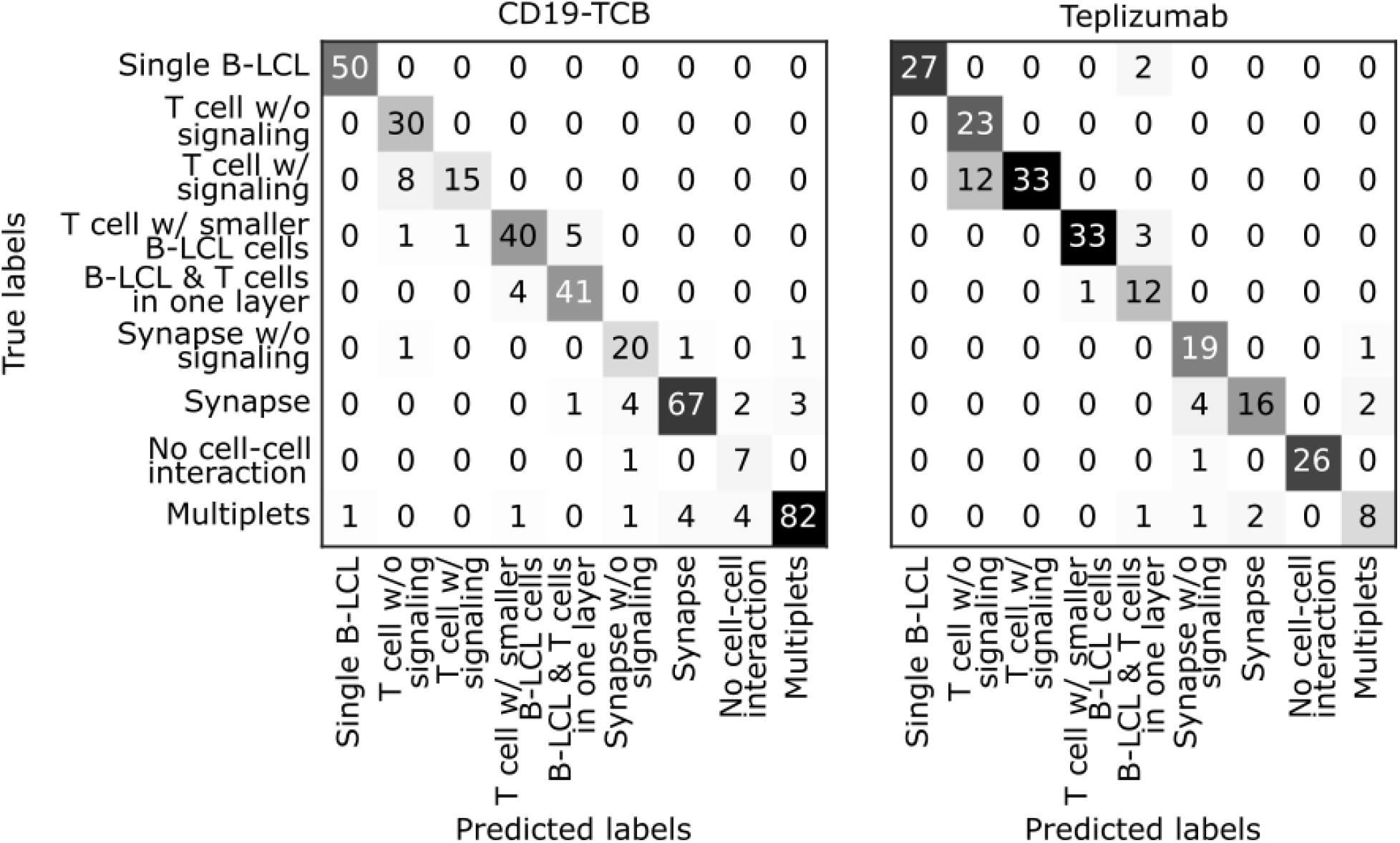
Inter-experiment and inter-stimulation Generalizability. Confusion matrix for classifications in CD19-TCB and Teplizumab based on 396 and 227 expert-annotated images, respectively. The previously trained model (Fig. 1c) reached a macro F1-score of 0.86 and 0.85, respectively, on both datasets.

**Supplementary Figure 8.**
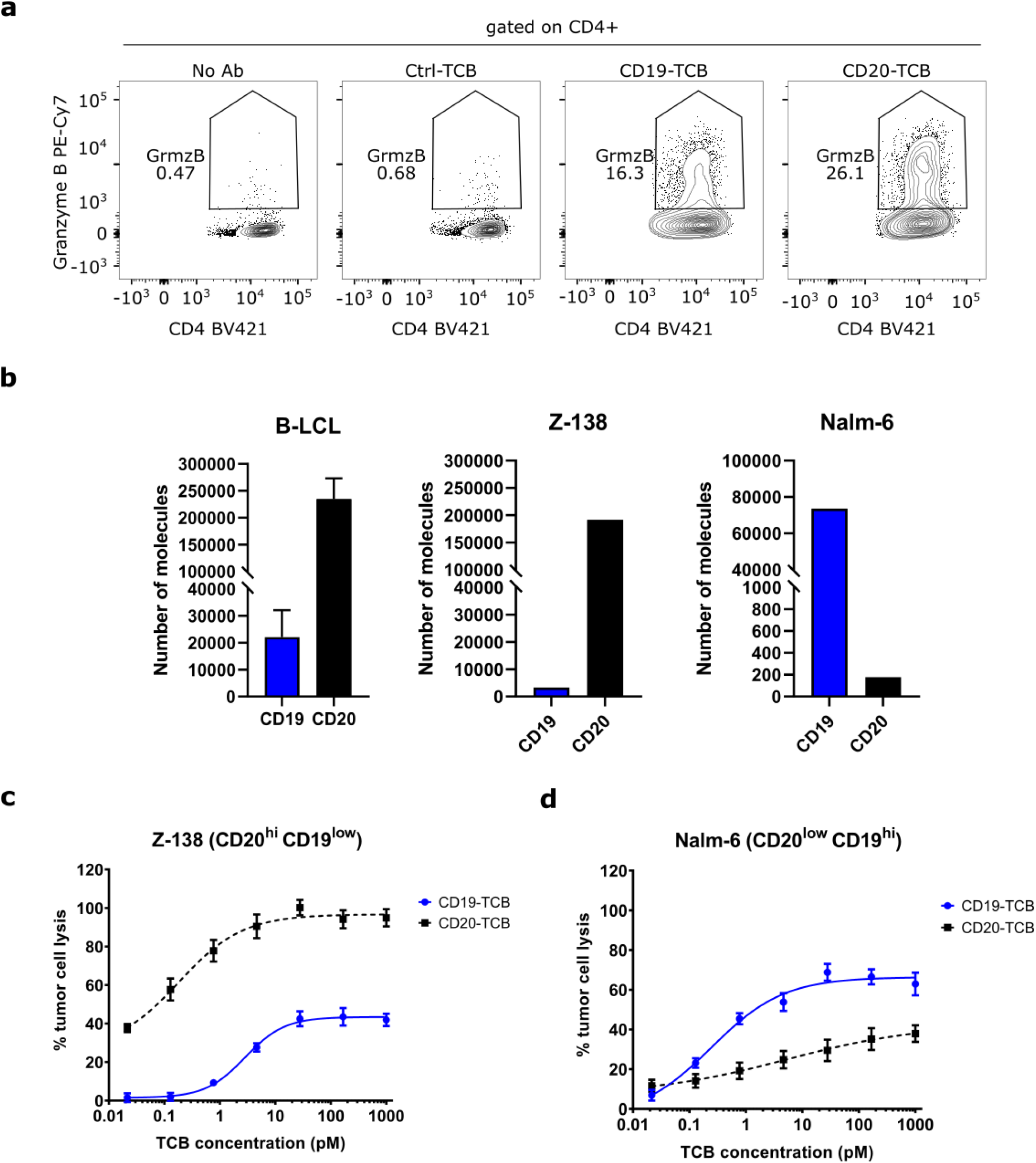
Investigation of GrzmB release and tumor killing induced by CD19- and CD20-TCB. **a** FACS plot showing GrzmB^+^ CD4^+^ T cells after treating the cells with Ctrl-TCB, CD19-TCB or CD20-TCB for 24h or left untreated (no Ab). A representative example of the gating strategy is shown in Supplementary Fig. 9a. **b** Number of CD19 and CD20 molecules expressed on B-LCL, Nalm-6 and Z-138 cells. The measured values are derived from cell culture and were not measured in parallel to the experiment. **c,d** Tumor cell lysis induced by CD19- and CD20-TCB, determined using LDH release after 24 h incubation of B-cell depleted human PBMCs with the tumor targets Nalm-6 or Z-138 and indicated TCB concentrations. One donor is shown.

**Supplementary Figure 9.**
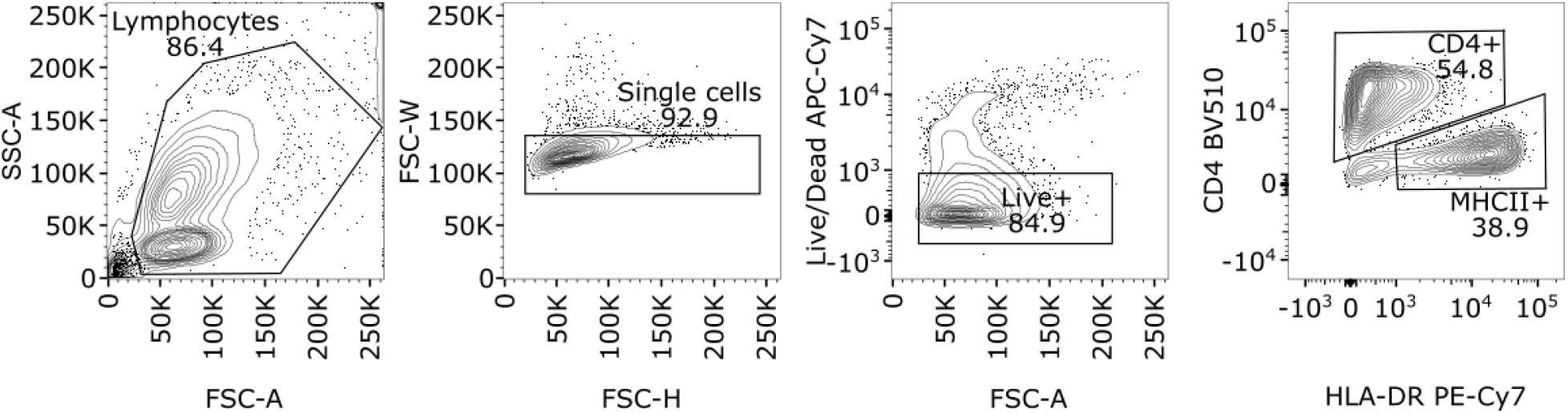
Gating strategy of conventional flow cytometry data. Representative example of the gating strategy used for analyzing conventional flow cytometry data as shown in Supplementary Fig. 1d and Supplementary Fig. 8a. Briefly, lymphocytes were selected in the FSC-A and SSC-A gate. In the next step, single cells were selected using FSC-H/FSC-W and viable cells were identified using the fixable viability dye eF780 (gated on eF780 negative cells). Finally, cells were gated on CD4^+^ T cells.

**Supplementary Table 1.**
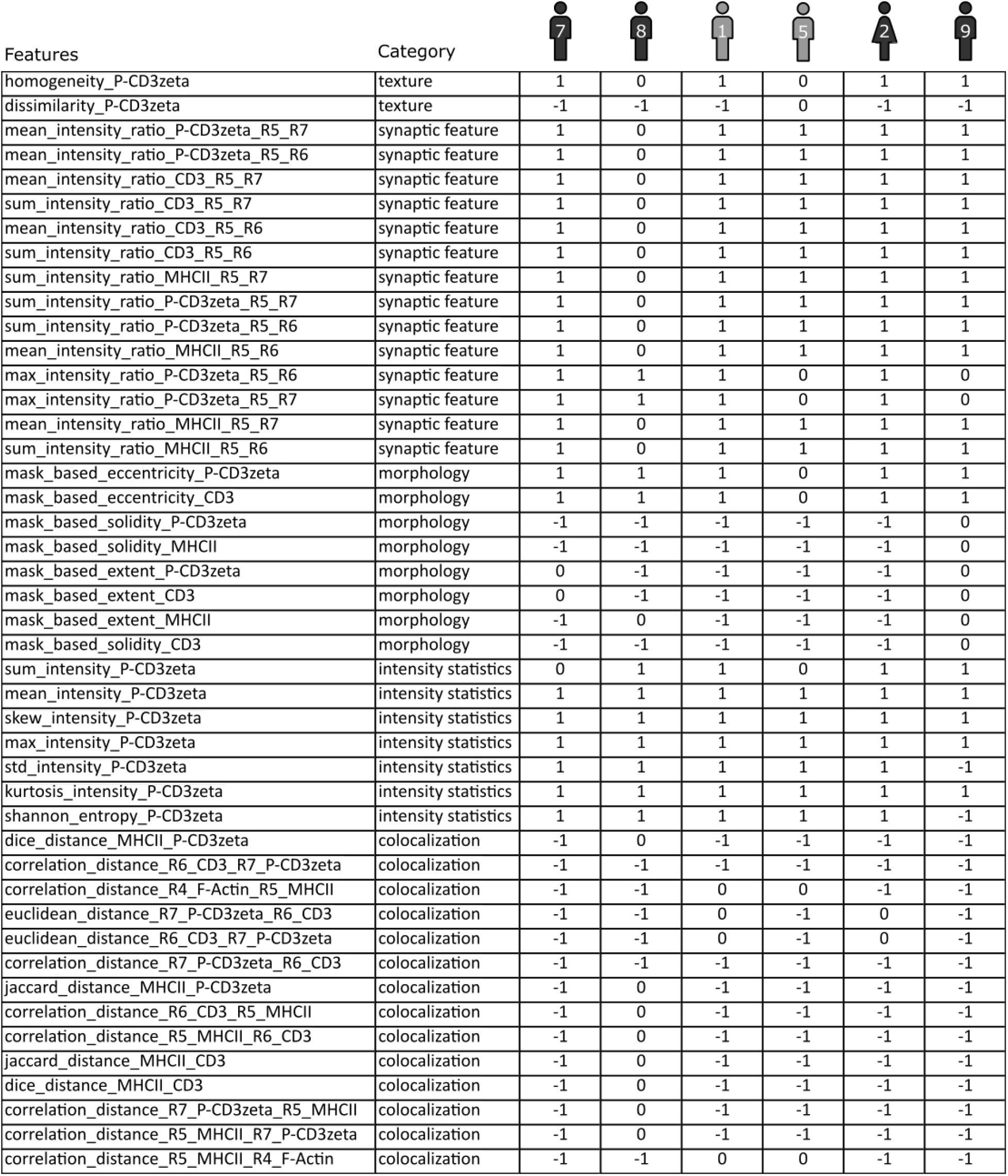
Significant features induced by CD19-TCB. The table represents the features that significantly changed for at least four donors due to stimulation by CD19-TCB. The table represents the list of consistent features from Fig. 3a. 1 represents a significant increase (red in Fig. 3a), -1 represents a significant decrease (blue in Fig. 3a) and 0 represents no significant change (gray in Fig. 3a).

| Features | Category |  |  |  |  |  |  |
| --- | --- | --- | --- | --- | --- | --- | --- |
|  |  | 7 | 8 | 1 | 5 | 2 | 9 |
| homogeneity_P-CD3zeta | texture | 1 | 0 | 1 | 0 | 1 | 1 |
| dissimilarity_P-CD3zeta | texture | -1 | -1 | -1 | 0 | -1 | -1 |
| mean_intensity_ratio_P-CD3zeta_R5_R7 | synaptic feature | 1 | 0 | 1 | 1 | 1 | 1 |
| mean_intensity_ratio_P-CD3zeta_R5_R6 | synaptic feature | 1 | 0 | 1 | 1 | 1 | 1 |
| mean_intensity_ratio_CD3_R5_R7 | synaptic feature | 1 | 0 | 1 | 1 | 1 | 1 |
| sum_intensity_ratio_CD3_R5_R7 | synaptic feature | 1 | 0 | 1 | 1 | 1 | 1 |
| mean_intensity_ratio_CD3_R5_R6 | synaptic feature | 1 | 0 | 1 | 1 | 1 | 1 |
| sum_intensity_ratio_CD3_R5_R6 | synaptic feature | 1 | 0 | 1 | 1 | 1 | 1 |
| sum_intensity_ratio_MHCII_R5_R7 | synaptic feature | 1 | 0 | 1 | 1 | 1 | 1 |
| sum_intensity_ratio_P-CD3zeta_R5_R7 | synaptic feature | 1 | 0 | 1 | 1 | 1 | 1 |
| sum_intensity_ratio_P-CD3zeta_R5_R6 | synaptic feature | 1 | 0 | 1 | 1 | 1 | 1 |
| mean_intensity_ratio_MHCII_R5_R6 | synaptic feature | 1 | 0 | 1 | 1 | 1 | 1 |
| max_intensity_ratio_P-CD3zeta_R5_R6 | synaptic feature | 1 | 1 | 1 | 0 | 1 | 0 |
| max_intensity_ratio_P-CD3zeta_R5_R7 | synaptic feature | 1 | 1 | 1 | 0 | 1 | 0 |
| mean_intensity_ratio_MHCII_R5_R7 | synaptic feature | 1 | 0 | 1 | 1 | 1 | 1 |
| sum_intensity_ratio_MHCII_R5_R6 | synaptic feature | 1 | 0 | 1 | 1 | 1 | 1 |
| mask_based_eccentricity_P-CD3zeta | morphology | 1 | 1 | 1 | 0 | 1 | 1 |
| mask_based_eccentricity_CD3 | morphology | 1 | 1 | 1 | 0 | 1 | 1 |
| mask_based_solidity_P-CD3zeta | morphology | -1 | -1 | -1 | -1 | -1 | 0 |
| mask_based_solidity_MHCII | morphology | -1 | -1 | -1 | -1 | -1 | 0 |
| mask_based_extent_P-CD3zeta | morphology | 0 | -1 | -1 | -1 | -1 | 0 |
| mask_based_extent_CD3 | morphology | 0 | -1 | -1 | -1 | -1 | 0 |
| mask_based_extent_MHCII | morphology | -1 | 0 | -1 | -1 | -1 | 0 |
| mask_based_solidity_CD3 | morphology | -1 | -1 | -1 | -1 | -1 | 0 |
| sum_intensity_P-CD3zeta | intensity statistics | 0 | 1 | 1 | 0 | 1 | 1 |
| mean_intensity_P-CD3zeta | intensity statistics | 1 | 1 | 1 | 1 | 1 | 1 |
| skew_intensity_P-CD3zeta | intensity statistics | 1 | 1 | 1 | 1 | 1 | 1 |
| max_intensity_P-CD3zeta | intensity statistics | 1 | 1 | 1 | 1 | 1 | 1 |
| std_intensity_P-CD3zeta | intensity statistics | 1 | 1 | 1 | 1 | 1 | -1 |
| kurtosis_intensity_P-CD3zeta | intensity statistics | 1 | 1 | 1 | 1 | 1 | 1 |
| shannon_entropy_P-CD3zeta | intensity statistics | 1 | 1 | 1 | 1 | 1 | -1 |
| dice_distance_MHCII_P-CD3zeta | colocalization | -1 | 0 | -1 | -1 | -1 | -1 |
| correlation_distance_R6_CD3_R7_P-CD3zeta | colocalization | -1 | -1 | -1 | -1 | -1 | -1 |
| correlation_distance_R4_F-Actin_R5_MHCII | colocalization | -1 | -1 | 0 | 0 | -1 | -1 |
| euclidean_distance_R7_P-CD3zeta_R6_CD3 | colocalization | -1 | -1 | 0 | -1 | 0 | -1 |
| euclidean_distance_R6_CD3_R7_P-CD3zeta | colocalization | -1 | -1 | 0 | -1 | 0 | -1 |
| correlation_distance_R7_P-CD3zeta_R6_CD3 | colocalization | -1 | -1 | -1 | -1 | -1 | -1 |
| jaccard_distance_MHCII_P-CD3zeta | colocalization | -1 | 0 | -1 | -1 | -1 | -1 |
| correlation_distance_R6_CD3_R5_MHCII | colocalization | -1 | 0 | -1 | -1 | -1 | -1 |
| correlation_distance_R5_MHCII_R6_CD3 | colocalization | -1 | 0 | -1 | -1 | -1 | -1 |
| jaccard_distance_MHCII_CD3 | colocalization | -1 | 0 | -1 | -1 | -1 | -1 |
| dice_distance_MHCII_CD3 | colocalization | -1 | 0 | -1 | -1 | -1 | -1 |
| correlation_distance_R7_P-CD3zeta_R5_MHCII | colocalization | -1 | 0 | -1 | -1 | -1 | -1 |
| correlation_distance_R5_MHCII_R7_P-CD3zeta | colocalization | -1 | 0 | -1 | -1 | -1 | -1 |
| correlation_distance_R5_MHCII_R4_F-Actin | colocalization | -1 | -1 | 0 | 0 | -1 | -1 |

**Supplementary Table 2.**
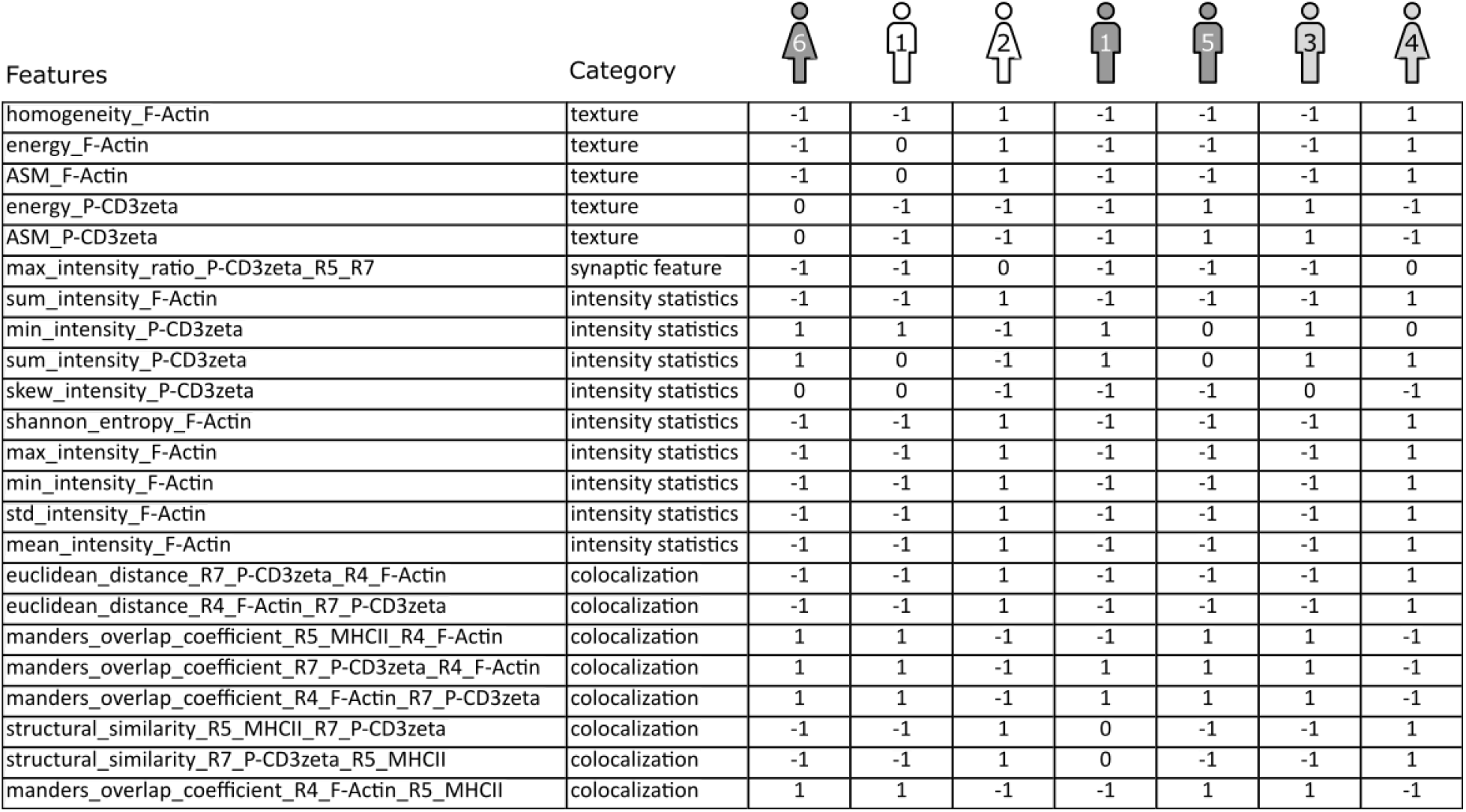
Significant features induced by Teplizumab. The table represents the features that significantly changed for at least six donors due to stimulation by Teplizumab. The table represents the list of consistent features from Fig. 3h. 1 represents a significant increase (red in Fig. 3h), -1 represents a significant decrease (blue in Fig. 3h) and 0 represents no significant change (gray in Fig. 3h).

| Features | Category |  |  |  |  |  |  |  |
| --- | --- | --- | --- | --- | --- | --- | --- | --- |
| homogeneity_F-Actin | texture | -1 | -1 | 1 | -1 | -1 | -1 | 1 |
| energy_F-Actin | texture | -1 | 0 | 1 | -1 | -1 | -1 | 1 |
| ASM_F-Actin | texture | -1 | 0 | 1 | -1 | -1 | -1 | 1 |
| energy_P-CD3zeta | texture | 0 | -1 | -1 | -1 | 1 | 1 | -1 |
| ASM_P-CD3zeta | texture | 0 | -1 | -1 | -1 | 1 | 1 | -1 |
| max_intensity_ratio_P-CD3zeta_R5_R7 | synaptic feature | -1 | -1 | 0 | -1 | -1 | -1 | 0 |
| sum_intensity_F-Actin | intensity statistics | -1 | -1 | 1 | -1 | -1 | -1 | 1 |
| min_intensity_P-CD3zeta | intensity statistics | 1 | 1 | -1 | 1 | 0 | 1 | 0 |
| sum_intensity_P-CD3zeta | intensity statistics | 1 | 0 | -1 | 1 | 0 | 1 | 1 |
| skew_intensity_P-CD3zeta | intensity statistics | 0 | 0 | -1 | -1 | -1 | 0 | -1 |
| shannon_entropy_F-Actin | intensity statistics | -1 | -1 | 1 | -1 | -1 | -1 | 1 |
| max_intensity_F-Actin | intensity statistics | -1 | -1 | 1 | -1 | -1 | -1 | 1 |
| min_intensity_F-Actin | intensity statistics | -1 | -1 | 1 | -1 | -1 | -1 | 1 |
| std_intensity_F-Actin | intensity statistics | -1 | -1 | 1 | -1 | -1 | -1 | 1 |
| mean_intensity_F-Actin | intensity statistics | -1 | -1 | 1 | -1 | -1 | -1 | 1 |
| euclidean_distance_R7_P-CD3zeta_R4_F-Actin | colocalization | -1 | -1 | 1 | -1 | -1 | -1 | 1 |
| euclidean_distance_R4_F-Actin_R7_P-CD3zeta | colocalization | -1 | -1 | 1 | -1 | -1 | -1 | 1 |
| manders_overlap_coefficient_R5_MHCII_R4_F-Actin | colocalization | 1 | 1 | -1 | -1 | 1 | 1 | -1 |
| manders_overlap_coefficient_R7_P-CD3zeta_R4_F-Actin | colocalization | 1 | 1 | -1 | 1 | 1 | 1 | -1 |
| manders_overlap_coefficient_R4_F-Actin_R7_P-CD3zeta | colocalization | 1 | 1 | -1 | 1 | 1 | 1 | -1 |
| structural_similarity_R5_MHCII_R7_P-CD3zeta | colocalization | -1 | -1 | 1 | 0 | -1 | -1 | 1 |
| structural_similarity_R7_P-CD3zeta_R5_MHCII | colocalization | -1 | -1 | 1 | 0 | -1 | -1 | 1 |
| manders_overlap_coefficient_R4_F-Actin_R5_MHCII | colocalization | 1 | 1 | -1 | -1 | 1 | 1 | -1 |

**Supplementary Table 3.** Glossary. The complete list of all the abbreviations in the text

|  |  |
| --- | --- |
| APC | Antigen-presenting cell |
| Brightfield | BF |
| EBV | Epstein-Barr virus |
| FACS | Fluorescence-activated cell sorting |
| GrzmB | Granzyme B |
| HPC | High Performance Computing |
| IFC | Imaging flow cytometry |
| MSE | Mean squared error |
| TCR | T cell receptor |
| MHC | Major histocompatibility complex |
| PBMCs | Peripheral blood mononuclear cells |
| RGB | Red Green Blue |
| Staphylococcus aureus enterotoxin A | SEA |
| TCB | T cell bispecific |
| UMAP | Uniform Manifold Approximation and Projection for Dimension Reduction |

